# Constraints on the xylem safety-efficiency trade-off

**DOI:** 10.64898/2026.09.25.754362

**Authors:** Fon Robinson Tezeh, Roman Mathias Link, Peter Petrik, Ronny Richter, Alexandra Weigelt, Christian Wirth, Bernhard Schuldt

## Abstract

- The xylem safety–efficiency trade-off is a central paradigm in plant hydraulics, yet empirical support remains inconsistent. In this study, we highlight how this inconsistency may arise from variation in environmental and evolutionary constraints and from differences in observational scale.
- We analysed 38 diffuse-porous angiosperm and 24 conifer species grown in a common garden experiment to quantify the relationship between embolism resistance (P50) and specific hydraulic conductivity (Ks) across and within species, and its differences between three species groups differing in evolutionary ancestry and eco-hydrological strategy.
- When pooling over all species, we found evidence for a safety-efficiency trade-off both across and within species. However, this pattern was driven almost exclusively by the mesic broadleaved species. In contrast, the across-species pattern was suppressed in coniferous species due to the evolutionary constraints on efficiency posed by their tracheid-based wood anatomy, while there was no evidence for a trade-off on either level in hydrophilic broadleaved species, which evolved under conditions that pose reduced selective pressure on embolism resistance.
- Our results demonstrate that the safety–efficiency trade-off is not universal, but emerges only in specific ecological and evolutionary contexts, highlighting the importance of multi-scale analyses and constraint-based frameworks for interpreting plant hydraulic strategies.

## Introduction

To maintain photosynthesis, plants have to efficiently transport water from the roots to the leaves. Under severe drought stress, this ability for water transport is impaired by the formation of xylem emboli, i.e. gas bubbles that expand and permanently block xylem conduits. While it would be advantageous for plants to maximize both safety against embolism and hydraulic efficiency, such trait combinations are exceedingly rare (Gleason et al., 2016). A long-standing tenet of plant hydraulics is therefore that hydraulic safety trades off with the hydraulic efficiency of the xylem, with plants having a more embolism resistant xylem facing a higher flow resistance when transporting water to their leaves and vice versa (Zimmermann, 1983). In this context, safety is usually defined as the (negative) water potential at 50 % loss of xylem hydraulic conductivity (−*P*_50_, MPa) and efficiency as the sapwood-specific hydraulic conductivity (*K*_s_, kg m^−1^ MPa^−1^ s^−1^). While the existence of this trade-off has long been taken for granted on theoretical grounds, the empirical evidence is mixed, with large-scale meta-studies only providing support for a relatively weak relationship between *P*_50_ and *K*_s_ across species (Gleason et al., 2016; Liu et al., 2021). Indeed, Gleason et al. (2016) identified the question why a large fraction of vascular species has both a low safety and a low efficiency, rendering them ‘incompetent’ (Liu et al., 2021) both in terms of water transport and of drought sensitivity, as central for understanding the evolution of xylem.

Several mechanisms have been identified as potential explanations for the existence of a safety-efficiency trade-off (Box 1). Given the inescapable nature of the proposed physical mechanisms, the weak relationship between *P*_50_ and *K*_s_ found in large datasets despite covering a considerable range in both traits (Gleason et al., 2016; Liu et al., 2021) seems surprising. One potential explanation for the weak link is that plants have many ways to modify efficiency by adjusting xylem structure that do not always affect safety (Gleason et al., 2016). Moreover, the trade-off may not be equally selected for in different climates and in plants following different ecological strategies (Gleason et al., 2016; Liu et al., 2021). Further, concomitant trade-offs between efficiency and mechanical strength (Bittencourt et al., 2016; Brodersen et al., 2016) and/or safety and storage (Pratt & Jacobsen, 2017) may obscure the trade-off, and the predominant focus on area-based metrics of efficiency may misrepresent its nature (Bittencourt et al., 2016). In addition, it has been pointed out that selection for trait combinations acts on the level of organisms rather than single tissues, and in consequence, it may be more promising to study trade-offs in an integrated approach considering traits throughout the entire plant instead of focusing solely on xylem traits (Meinzer et al., 2010).

To better understand the safety-efficiency trade-off, some theoretical background on the concept of evolutionary trade-offs is beneficial. From a theoretical perspective, two traits are considered to *trade off* with each other when “an increase in fitness due to a change in one trait is opposed by a decrease in fitness due to a concomitant change in the second trait” (Roff & Fairbairn, 2007). Many types of trade-offs are assumed to arise because traits are subject to different types of *evolutionary constraints*, i.e., “restrictions or limitations on the course or outcome of evolution” (Arnold, 1992). The mechanisms for the safety-efficiency trade-off in Box 1 largely result from *functional constraints* (Garland et al., 2022) at the pit membrane, conduit and/or tissue level. Functional constraints arise when features that enhance the performance of one task (e.g., a thicker pit membrane limiting the passage of bubbles and/or dissolved gas) decrease the performance of another (e.g., reduced hydraulic conductance) (Holzman et al., 2011; Shoval et al., 2012). At the tissue level, these functional constraints can be exacerbated by *allocation constraints* (Garland et al., 2022), which arise from the limited space in the xylem (Box 1), particularly when additional trade-offs with other wood properties are taken into account (Bittencourt et al., 2016; Brodersen et al., 2016; Pratt & Jacobsen, 2017).

It is important to point out that evolutionary constraints, as well as the resulting trade-offs, can be context-dependent (Laitinen & Nikoloski, 2024). This is particularly true for trade-offs resulting from constraints imposed by the environment rather than e.g. from functional constraints or the allocation of limited resources (Garland et al., 2022). The existence of such a trade-off does not automatically imply observable negative trait correlations within or between species – ecological circumstances define whether there is a selective regime (Garland et al., 2022) that eliminates less favourable trait combinations. If driven by such processes, the safety-efficiency trade-off might not affect all species equally. This is supported by Liu et al. (2021), who noted that species from drier and more seasonal climates were more likely to co-optimize safety and efficiency than species from wetter habitats, and that conservative water use strategies as well as a reliance on ample water storage may release species from high safety requirements.

Whether or not an existing trade-off between traits results in an observable negative trait correlation moreover depends to a great extent on the scale of aggregation the data are analysed on, as different processes matter on individual and on species level. If, as generally expected, the safety-efficiency trade-off resulted from functional or allocation constraints resulting from xylem anatomy, the strongest relationships would be expected on the intraspecific level, but would not necessarily translate into patterns on higher levels (Agrawal, 2020). The reason for this expectation is that while these constraints impose physical limits on how two traits can covary, across species these patterns may be confounded due to variation in these traits resulting from other sources. If, however, selection and niche specialization in response to the environment were the main driving forces of the trade-off, we would expect patterns to emerge primarily across species (Agrawal, 2020). While the importance of scale for the interpretation of the safety-efficiency trade-off has often been pointed out (Sperry et al., 2008; Meinzer et al., 2010), so far, the most compelling evidence is based on interspecies patterns (Gleason et al., 2016; Liu et al., 2021), and there are no principled multi-level analyses of that investigate the relationship between *P*_50_ and *K*_s_ on multiple scales of aggregation.

In this study, we analysed 62 tree species grown together in a common garden experiment to study the safety-efficiency trade-off within and across species under controlled conditions that limit the effects of age and environment on the analysed traits. We focused on three species groups defined by evolutionary ancestry and eco-hydrological strategy: conifers, hydrophilic broad-leaved trees and mesic broad-leaved trees. We hypothesized i) that safety (−*P*_50_) and efficiency (*K*_s_) are negatively correlated both within and across species, and ii) that these correlations differ between our species groups showing: (a) the strongest correlations in mesic broad-leaved tree species, (b) suppressed interspecific correlations in conifers due to the strong evolutionary constraints on conduit anatomy that limit the range of their hydraulic efficiency, and (c) no or weaker patterns in hydrophilic tree species, which evolved in environments with a reduced selective pressure on hydraulic safety. We further tested whether iii) within-species patterns are stronger than across-species patterns, as would be expected if these patterns resulted from functional constraints on the pit membrane, conduit or tissue level.

## Materials and methods

### Study site and plant material

For this study, we used data from branch samples from 236 trees from 38 diffuse-porous angiosperm species and 24 coniferous tree species from 31 genera and 14 families (Table S1) from the ARBOfun research arboretum near Leipzig, Germany (51°16′N, 12°30′E; 150 m a.s.l.). Established between 2012 and 2014, the 2.5-hectare arboretum hosts 100 tree species from 39 families originating from hemi-boreal to sub-mediterranean forests in Europe, as well as selected species from Asia and North America (Kretz et al., 2025). The common garden design allowed us to minimise environmental and ontogenetic variation and isolate intrinsic interspecific differences in hydraulic traits. For the study, all trees in the arboretum were targeted that were big enough to permit sample excision, with the exception of ring-porous and semi-ring-porous species, which cannot be reliably measured with the flow centrifuge method (Cochard et al., 2010; Torres-Ruíz et al., 2014). The trees were planted 5.8 m apart in a randomized block design with five blocks, each of which containing one individual of each species. The region experiences a mean annual precipitation of approximately 520 mm and a mean annual temperature of 9.7 °C (1980–2020; DWD Climate Data Center [CDC], Station Leipzig/Halle, ID 2932). The soil of the arboretum, formerly part of a managed arable field, is classified as a Luvisol with a pH of 5.7 (Ferlian et al., 2017).

Plant material from the arboretum was collected between May and September 2023 from trees aged ca. 11 years (Table S1). Samples were collected from trees in all blocks from all healthy individuals of the target species suitable for hydraulic measurements (i.e., having enough material to permit taking at least one sufficiently sized sample). At the arboretum, 40 to 70 cm long twigs were cut from the uppermost sun-exposed canopy from either the east or west-facing side, wrapped in wet paper towels, bagged in dark humidified plastic bags, and transported to the laboratory, where they were stored in a cooling chamber at 5 °C and processed within two weeks (cf. Herbette et al., 2010).

### Species groups

Due to the pronounced differences in wood anatomy, the studied species were separated into conifers (24 spp.) and broadleaved trees (38 spp.). To explain the extreme differences in behaviour among the broadleaved species, the latter were further separated into two subgroups, namely mesic (23 spp.) and hydrophilic broadleaved species (15 spp.; Table S1). The classification of the “hydrophilic” angiosperm species was based on expert knowledge and literature, and refers to species adapted to high-humidity habitats such as softwood floodplain forests, swamps and bogs (Table S2). As riparian species in particular are adapted to a high disturbance regime (Tabacchi et al., 1998), this group strongly overlaps in their adaptations with pioneer species, and some species that predominantly grow as pioneers (*Alnus spp.*, *Salix spp.*) but reliably occur in riverine habitats were also included in the group of hydrophilic species. All remaining angiosperm species were assigned as “mesic”.

### Ellenberg-Tichý indicator values

To illustrate differences in habitat preferences between the three species groups, we obtained Ellenberg-Tichý indicator values for a subset of the studied species based on Tichý et al. (2023), a recent synthesis of Ellenberg-type indicator values (EIV) for European plant species. From this source, we extracted the indicator values for moisture, temperature and light, obtaining values for 27, 28 and 30 species, respectively. The EIV for moisture ranges describes plant moisture preferences, with higher values denoting species with a preference for moister habitats (Ellenberg et al., 2001). The EIV for temperature describes the geographical and altitudinal range of species with lower values in alpine and montane habitats and higher values denoting species occurring in warmer and increasingly more Mediterranean habitats (Ellenberg et al., 2001). The EIV for light describes the preferences of a species on a spectrum from deep shade to full sunlight, with higher values representing species that occur in more open habitats (Ellenberg et al., 2001).

### Quantification of xylem hydraulic conductivity

To determine branch xylem hydraulic conductivity (*K*_h_; kg m s^-1^ MPa^-1^), the 236 branches were rehydrated in water for ca. 30 min in the laboratory after recutting under water several times using pruning shears to release potential tension in the xylem (Torres-Ruiz et al., 2015). For the broadleaved trees, fresh samples were cut to standardized segments of ca. 29 cm, lateral twigs were excised, and cut ends were sealed with a rapid-curing adhesive (Loctite 431 with activator 7452; Henkel, Düsseldorf, Germany) to minimize leakage. Segments of the broadleaved species were then connected via silicone tubing to a Xyl’em Plus embolism meter (Bronkhorst, Montigny-Les-Cormeilles, France) for stem hydraulic conductivity measurements. The system was perfused with a degassed, demineralized measurement solution containing 10 mM KCl and 1 mM CaCl_2_. After measuring initial hydraulic conductivity at a pressure head of 6 kPa for 5 min, samples were repeatedly flushed at a high pressure of 120 kPa for 10 min each to remove emboli. When the measured conductivity values did not change between subsequent flushes, we assumed that maximum hydraulic conductivity (*K*_h_^max^) was reached. For the conifers, straight segments without side twigs of a length of ca. 5 cm were excised under water, debarked and degassed in the aforementioned measurement solution for 48 h in a vacuum chamber to remove emboli, as flushing at high pressure in conifers may lead to reductions of hydraulic conductivity due to pit aspiration. The branches were removed from the solution on the day of measurement. The hydraulic conductivity after degassing, assumed to be equal to *K*_h_^max^, was measured under a pressure head of 6 kPa. For both groups of samples, hydraulic conductivity was calculated as *K*_h_ = *f ⋅ L* / Δ*P*, where *f* (kg s⁻^1^) is the flow rate, and Δ*P* (MPa) is the pressure drop along the length of the segment (*L*, m).

Specific hydraulic conductivity (*K*_s_; *K*g m^-1^ MPa^-1^ s^-1^) was then derived by dividing *K*_h_^max^ by the basipetal cross-sectional xylem area (*A*_cross_, m^2^) excluding bark.

### Vulnerability curves with the flow-centrifuge method

Vulnerability curves based on the flow-centrifuge method were measured with a Cavitron device (Cochard et al., 2005) built from a Sorval RC 5 Plus series centrifuge with manual control of rotation speed, and using the Cavisoft software (Cavisoft v.5.2.1, University of Bordeaux, Bordeaux, France). For the broadleaved species, vulnerability curves were measured on the branch segments from the *K*_s_ measurement after shortening them to a length of 27.5 cm (mean diameter at basipetal end ± SD: 6.87 ± 0.76 mm). For the conifers, the same segment length was excised from the branch from which the 5 cm for *K*_s_ measurement was cut, and lateral twigs excised and glued as described above. The conifer segments were then completely debarked to prevent resin blockage, while for the angiosperms, a 3 cm section of bark was removed from both ends to prevent water absorption under the bark. Sample diameters were measured twice at both sample ends. Using the measurement solution described above, we repeatedly measured conductance while progressively lowering the water potential by increasing rotational velocity, beginning at −0.834 MPa and continuing until we observed at least a 90% loss of the initial conductance. After each change in rotational velocity, we waited for two minutes before taking new conductance readings to permit the system to reach a new steady state.

To avoid vessel-length related measurement artefacts, we ensured that the vulnerability curves were s-shaped for all analysed species (Fig. S1, S2, S3). Further, for all samples, we recorded the distance from the basipetal sample end to the tip of the branch. While there was a positive correlation between distance to tip and *P*_50_, this was entirely driven by species differences, with hydrophilic species tending to have longer and narrower branches and less negative *P*_50_ values. After accounting for species differences, there was no evidence of a positive within-species correlation between distance to tip and *P*_50_.

**Figure 1.**
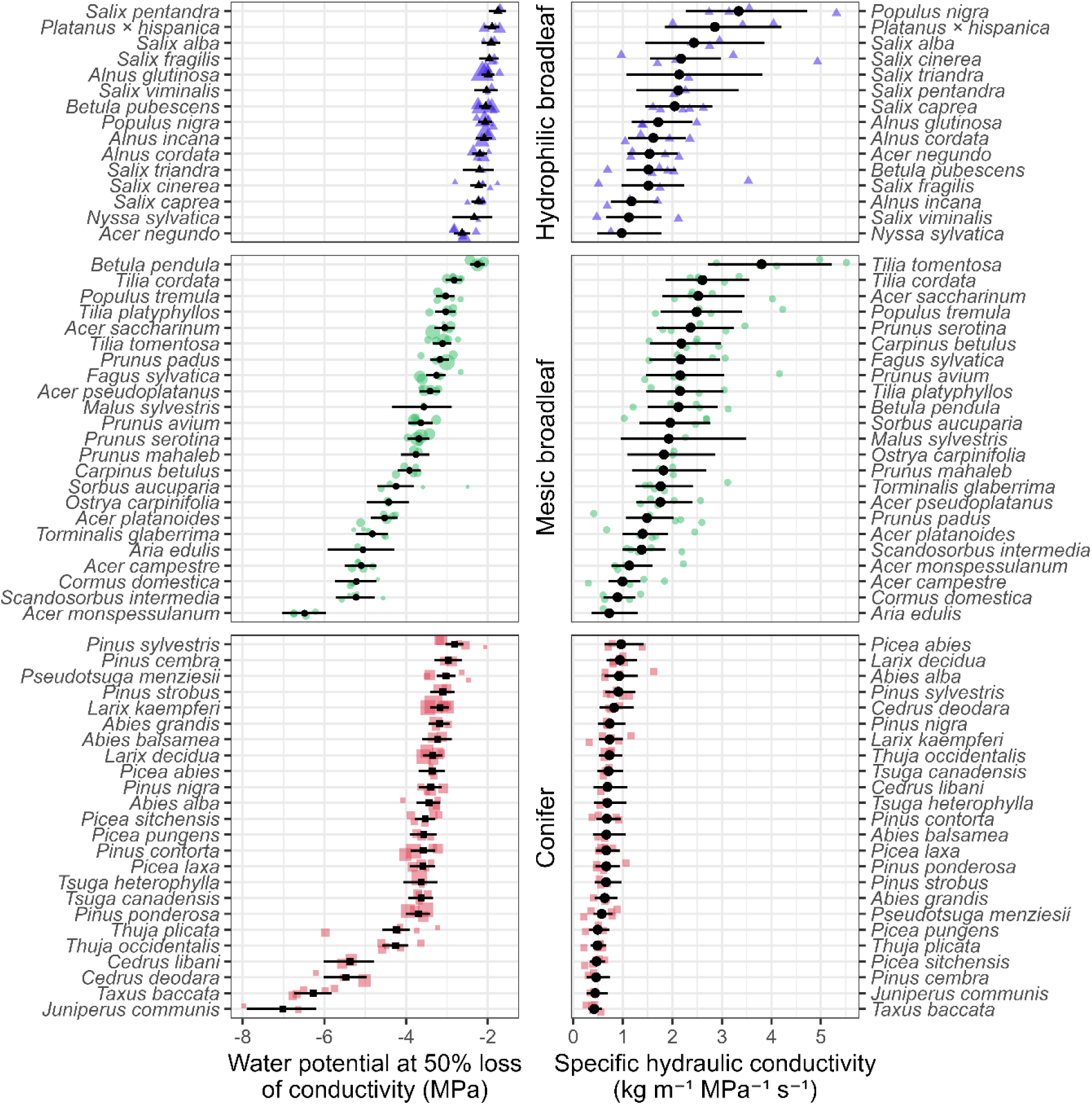
Observed ranges of xylem safety (*P*_50_; left) and hydraulic efficiency (*K*_s_; *K*_s_ (right) across 62 temperate tree species. The species are grouped into hydrophilic broadleaf species (n = 15), mesic broadleaf species (n = 23) and conifer species (n = 24). Shown are the raw data overlaid with the estimated posterior means and 95 % credible intervals from the model in Eqn. (2), (3). For *P*_50_, the size of the plotting symbol represents the precision (1 / SE^2^) of the *P*_50_ estimates, with more precise estimates displayed larger.

**Figure 2.**
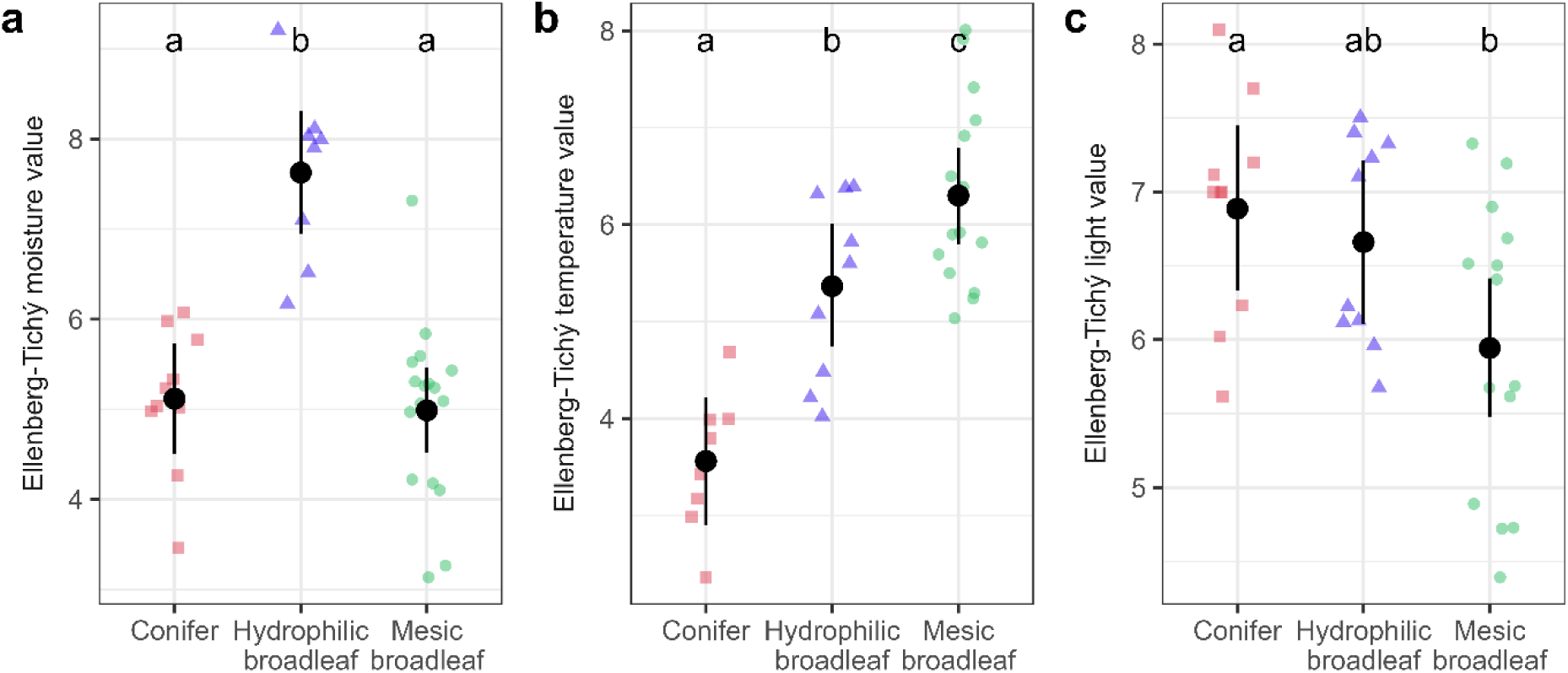
Ellenberg-Tichý indicator values of the different species groups. Shown are the raw data overlaid with the posterior means and 95 % credible intervals for a) the moisture value, b) the temperature value and c) the light value for conifers, mesic broadleaf and hydrophilic broadleaf species, respectively. Letters indicate credible differences between groups on the 0.05 level based on Bayesian ANOVA.

**Figure 3.**
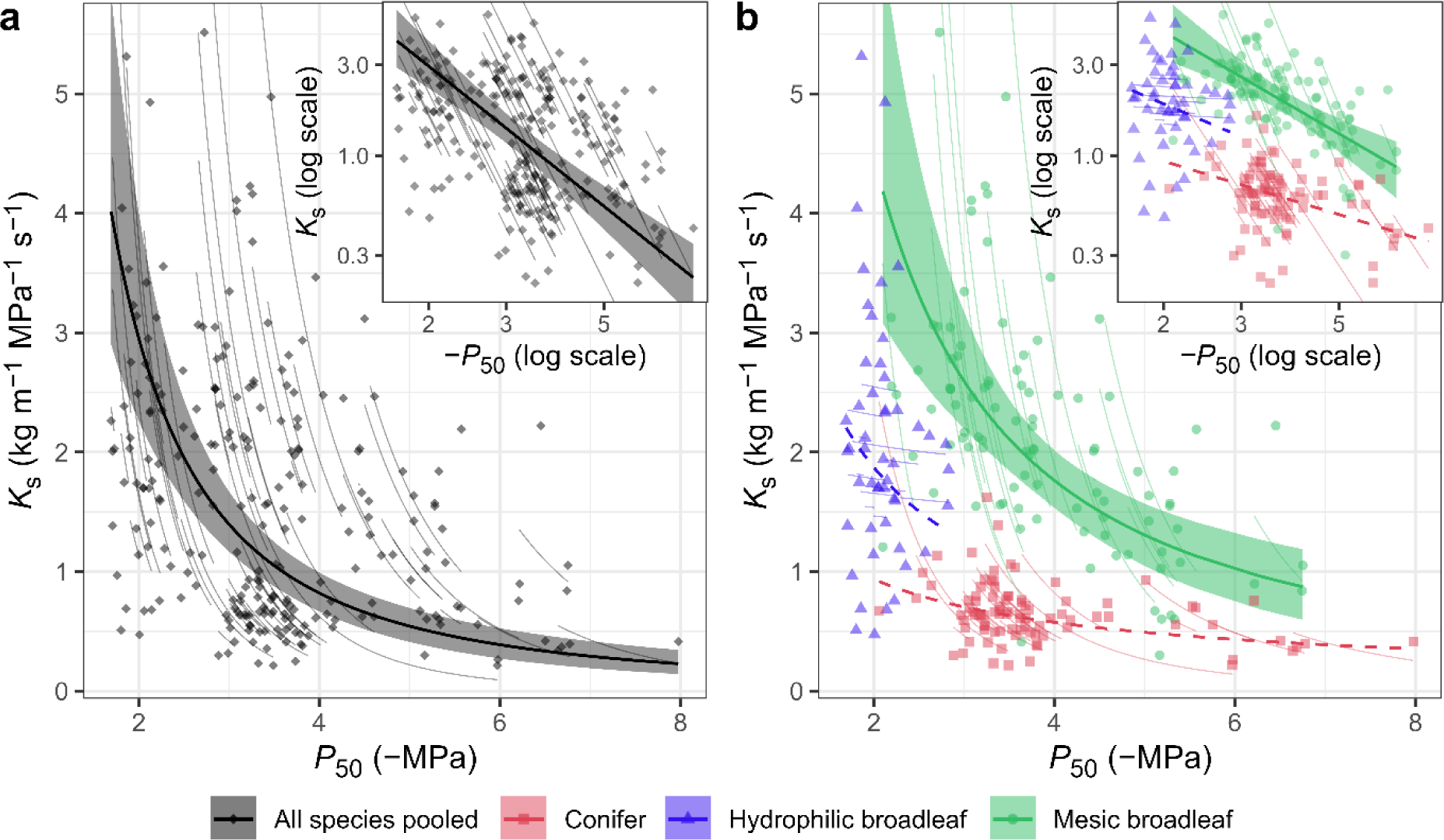
Model predictions for the relationship between xylem safety (*P*_50_) and hydraulic efficiency (*K*_s_). Observed data are shown together with the SMA regression lines for across-species (lines with 95 % credible intervals) and within-species relationships (thin lines) for a) the joint model across all species and b) separate models for each species group. Dashed lines indicate slopes with shaded 95 % credible intervals that do not exclude 0 (in which case no credible intervals are displayed, cf. Notes S1.4).

### Statistical analyses

All data handling and statistical analyses were performed in `R` v. 4.5.2 (R Core Team, 2025) in the framework of the `tidyverse` (Wickham et al., 2019).

Vulnerability curves were fitted in R v. 4.5.2 (R Core Team, 2025) with nonlinear least squares using a sigmoidal model (Pammenter and Vander Willigen, 1998) based on hydraulic conductivity (Ogle et al., 2009):

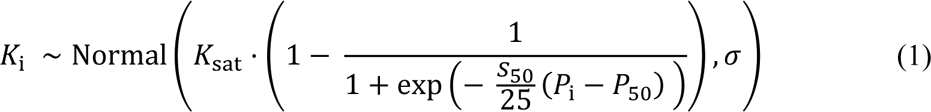

Here, for each observation *i*, the conductivity *K*_i_ is assumed to be normally distributed with residual standard deviation *σ* around a logistic function of the water potential *P*_i_ with the parameters *P*_50_ (water potential at 50% loss of conductivity), *S*_50_ (slope of the vulnerability curve on the percentage loss of conductivity scale) and *K*_sat_ (conductivity at full saturation).

For statistical testing, we used a Bayesian framework based on `R` package `brms` v.2.23.0 (Bürkner 2017; 2018). To test for differences in Ellenberg-Tichý indicator values between species groups, we performed Bayesian ANOVA using `brms` default priors. To assess the magnitude of the safety-efficiency trade-off within and across species, we described the relationship between safety (−*P*_50_) and efficiency (*K*_s_) in a mixed modelling framework that allowed us to separate the covariance between the two variables into across-and within-species components. To achieve this, we modelled natural log-transformed negative *P*_50_ (*_x_*_[*ij*]_) and natural log-transformed *K*_s_ (*y*_[*ij*]_) for observation *i* of *I* belonging to species *j* of *J* jointly in a multi-response mixed model with correlated residuals and correlated random species intercepts. The analysis was performed on the log-log-scale to account for the nonlinear nature of the relationship (cf. Pereira et al., 2023) and permit comparison with estimates in Gleason et al. (2016) and Liu et al. (2021). To account for the estimation uncertainty in the *P*_50_ estimates from Eqn. (1), we used a measurement error model for *_x_*_[*ij*]_:

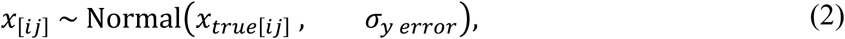

where *x_true_*_[*ij*]_ is the unmeasured true value of *x*_[*ij*]_ and *σ_error_* is the estimation uncertainty in log(−*P*_50_). *x_true_*_[*ij*]_ and *y*_[*ij*]_ where then jointly modeled as:

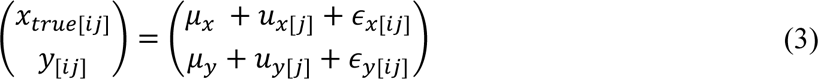

Here, *μ_x_* and *μ_y_* are the grand means, *u_x_*_[*j*]_ and *u_y_*_[*j*]_ are species-level random intercepts and *ε_x_*_[*ij*]_ and *ε_y_*_[*ij*]_ are the individual-level residuals for *x_true_*_[*ij*]_ and *y*_[*ij*]_, respectively. The species level random effects were assumed to follow a multivariate normal distribution with mean 0 and covariance matrix Σ*_across_*, while the residuals were modelled analogously with a multivariate normal distribution with mean 0 and covariance matrix Σ_*within*_:

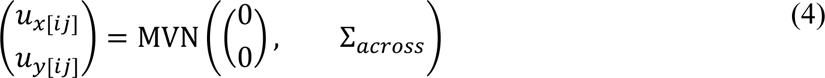

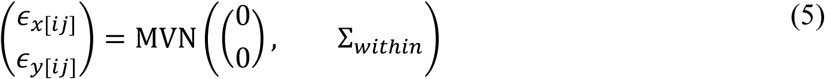

The across-species covariance matrix Σ*_across_* contains the across-species standard deviations

*τ_x_* and *τ_y_* as well as the across-species correlation *ρ_across_*, while the within-species covariance matrix Σ*_wit_*_ℎ*in*_ contains the within-species (residual) standard deviations *σ_x_* and *σ_y_* as well as the within-species correlation *ρ_wit_*_ℎ*in*_. Based on the estimated parameter values for within-and across-species correlations and standard deviations, we then calculated standardized major axis (SMA) regression type slopes and intercepts (Warton et al., 2006).

To test whether the trade-off differs between species groups, we fitted one model for all species as well as three separate models for hydrophilic broadleaf species, mesic broadleaf species and conifers. All models were fitted using moderately informative priors. Sampling was performed on 4 chains for 10000 iterations each, 5000 of which were discarded as warm-up. Convergence was evaluated based on the potential scale reduction factor *R̂* (Vehtari et al., 2021) and inspection of trace plots. Model adequacy was tested based on plots of posterior predictive checks (Gabry et al., 2019), leave-one-out cross-validation based on `R` package `loo` v. 2.9.0 (Vehtari et al., 2017, 2025) and residual diagnostics based on `R` package `DHARMa` v.0.4.7 (Hartig, 2024) via `DHARMA.helpers` v.0.0.2 (Rodríguez-Sánchez, 2026). The comparison between the two models was based on the expected log pointwise predictive density (ELPD), calculating the difference in ELPD between the joint and the separate models manually based on Vehtari et al. (2017). Parameter estimates were assumed to be credibly different from zero if their 95 % credible intervals excluded zero. Details about model structure, prior specifications and derived quantities can be found in Notes S1.

## Results

### Observed values of xylem safety and efficiency

Across the 62 analysed temperate tree species, we observed a wide range in water potential at 50 % loss of conductivity (*P*_50_), ranging from −1.70 MPa to −6.75 MPa for angiosperm species and −2.06 MPa to −7.98 MPa for conifer species (Figure 1, Table S1). For the angiosperms, clear habitat-related groupings emerged, with hydrophilic broadleaf species consistently exhibiting less negative *P*_50_ values (posterior mean −2.09 MPa; 95 % credible interval −2.20 – −1.98 MPa; Table S2, S4) than mesic broadleaves (–3.97 MPa, 95 CI: −4.38 – −3.60 MPa; Table S4, S5), indicating lower resistance to drought-induced embolism and hence lower hydraulic safety. With an average of −3.91 MPa (95 % CI: −4.31 – −3.55 MPa; Table S4), conifer species did not differ credibly from the mesic broadleaves (Table S5).

For sapwood-specific hydraulic conductivity (*K*_s_), angiosperms ranged from 0.30 to 5.51 kg m⁻¹ MPa⁻¹ s⁻¹, while conifers covered a much narrower range of 0.22 to 1.62 kg m^−^¹ MPa^−^¹ s^−^¹. Notably, we did not observe an average difference in *K*_s_ between mesic (2.17 kg m^−1^ MPa^−1^ s^−1^; 95 % CI: 1.86 – 2.55 kg m^−1^ MPa^−1^ s^−1^; Table S4) and hydrophilic broadleaf species (2.15 kg m^−1^ MPa^−1^ s^−1^; 95 % CI: 1.77 – 2.65 kg m^−1^ MPa^−1^ s^−1^; Table S4), while conifers had a credibly lower average of 0.66 kg m^−^¹ MPa^−^¹ s^−^¹ (95 % CI: 0.59 – 0.74 kg m^−1^ MPa^−1^ s^−1^; Table S4, S5).

### Differences in Ellenberg-Tichý indicator values

We found evidence for credible differences between species groups for all analysed Ellenberg-Tichý indicator values (Fig. 2, Table S6). The hydrophilic broadleaf species had moisture values that were on average 2.5 (95 % credible interval: 1.6 – 3.4) units higher than those for conifer species and on average 2.6 (1.8 – 3.5) units higher than those for mesic broadleaf species, confirming the preference for much wetter habitats of hydrophilic species (Fig. 2, Tables S6, S7). All three groups differed credibly from each other in their temperature values, with conifers on average having the lowest, hydrophilic broadleaf species intermediate and mesic broadleaf species the highest values, underlining the predominance of conifers in colder and more montane biomes (Fig. 2, Tables S6, S7). For the light values, conifers did not differ from hydrophilic broadleaf species, but both of the latter had on average higher values than mesic broadleaf species, indicating a preference of the latter for shadier habitats, at least during the seedling stage (Fig. 2, Tables S6, S7).

### Models of the safety-efficiency trade-off

When pooling over all species, we found a credible negative overall across-species relationship (ρ = −0.406, 95 % CI: −0.610 – −0.171) between specific hydraulic conductivity (log(*K*ₛ)) and hydraulic safety (log(−*P*_50_)), consistent with a trade-off between safety and efficiency (Table 1, Fig. 3a, Fig. S4, Table S3). Within species, there was a credible but weaker correlation between log(*K*_s_) and log(−*P*_50_) than across species (ρ = −0.269, 95 % CI: −0.415 – −0.111), though the within-species relationship had a credibly steeper slope (Fig. S4, Table S3, S8). The model explained a total of 71.8 % (95 % CI: 66.8 – 75.7 %) of the variance in log(*K*_s_) and 91.6 % (95 % CI: 90.4 – 92.5 %) of the variance in log(−*P*_50_) (Fig. 4), reflecting the higher within-species variability in *K*_s_.

**Figure 4.**
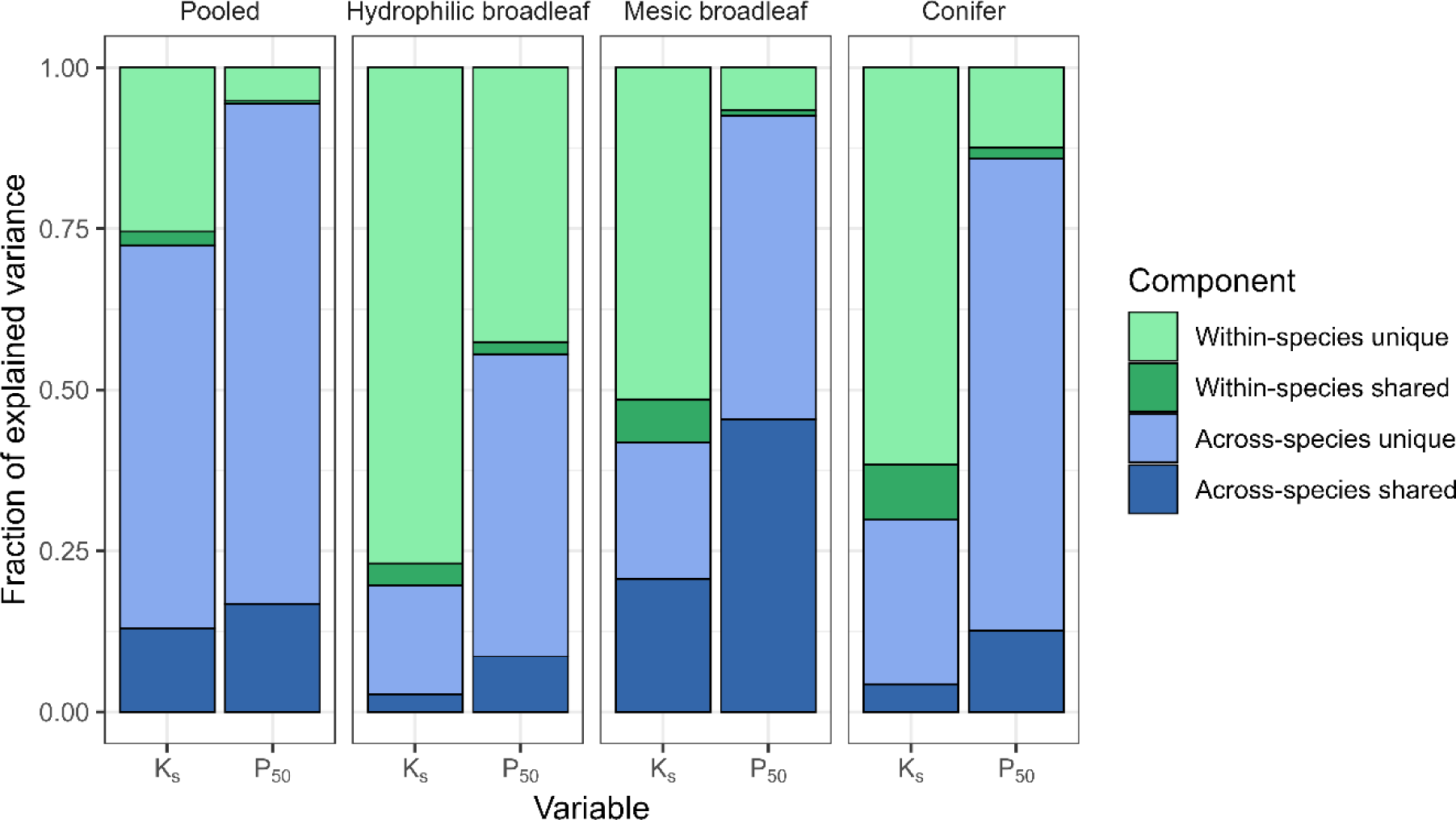
Variance decomposition for the pooled model and the three species-group-wise models. Shown are the posterior means of the fraction of variance explained by within-and across-species differences separated into the shared variance between *P*_50_ and *K*_s_ and the variance unique to each variable. Variance components were computed according to Eqn. (S19), (S20) in Notes S1.5.

**Table 1.**
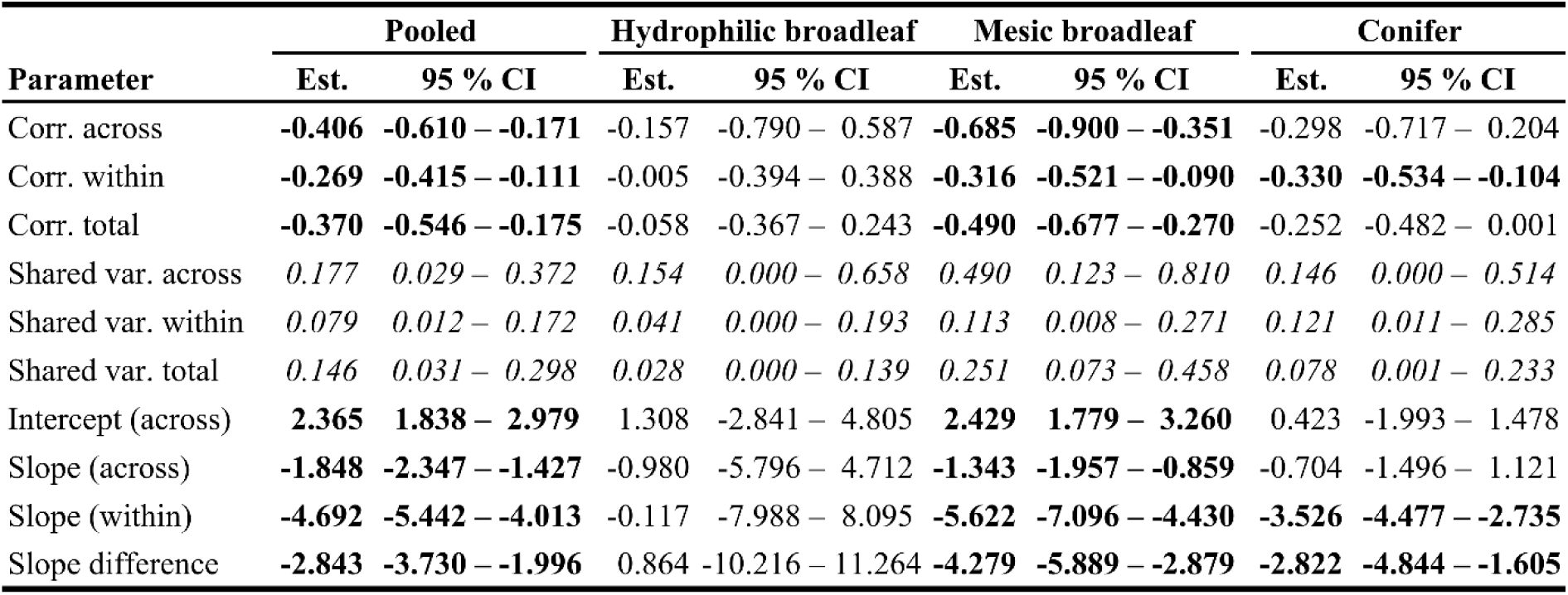
Summary of the key model parameters. For all four models (pooled over all species vs. separate models for hydrophilic broadleaf species, mesic broadleaf species and conifer species), the estimates of the correlation within and across species, the total correlation across organisational levels, the shared variance (R^2^ of the linear relationship) on the respective levels, the SMA intercept and slope as well as the difference between slopes on across-and within-species level are shown with their 95 % credible intervals. Parameters credibly different from zero on the 95% level are typeset in bold (shared variances are typeset in italics as they are constrained to values above zero). See Notes S1 for the calculation of total correlation and SMA parameters.

If separating by species groups, a different pattern emerged, with a much stronger negative across-species relationship between log(*K*_s_) and log(−*P*_50_) for mesic broadleaf species (ρ = −0.685, 95 % CI: −0.900 – −0.351) than in the pooled model, but no credible across-species correlation for conifers (ρ = −0.298, 95 % CI: −0.717 – 0.204) and hydrophilic broadleaf species (ρ = −0.157, 95 % CI: −0.790 – 0.587; Table 1, Fig. 3b, Fig. S4, Table S3). Within species, there was credible evidence for a negative safety-efficiency relationship in both mesic broadleaf trees (ρ = −0.316, 95 % CI: −0.521 – −0.090) and conifers (ρ = −0.330, 95 % CI: −0.534 – −0.104), both with steeper slopes than across species (Table 1, Fig. S4, Table S8). For hydrophilic broadleaf species, there neither was evidence for a within-species relationship (ρ = −0.005, 95 % CI: −0.394 – 0.388, Fig. S4, Table S3). The separate models on average explained a lower fraction of variance in log(*K*_s_) and log(−*P*_50_) (Fig. 4) due to the large fraction of variance explained by the differences among species groups. The hydrophilic broadleaf species had the lowest fraction of explained variance in both response variables (Fig. 4), showing the particularly high within-species variability in that group. The largest fraction of across-species variance shared between the response variables was found for the mesic broadleaf species (Fig. 4).

The model comparison based on leave-one-out cross-validation favoured the models with group-specific parameters by a ΔELPD of 14.9 (standard error: 9.9), indicating moderate evidence against the pooled model.

## Discussion

In our dataset comprising 38 diffuse-porous angiosperm and 24 conifer species, we found clear evidence for a negative relationship between (absolute) water potential at 50 % loss of conductivity (−*P*_50_) and specific hydraulic conductivity (*K*_s_), consistent with a safety-efficiency trade-off. This pattern was evident across scales of aggregation, although with a different slope across and within species. If analysed separately for species groups differing in evolutionary ancestry and eco-hydrological strategy, it became apparent that while the pattern was present both across and within species for mesic broadleaf species, it could only be found within species for conifers, and there was no evidence for a relationship on either observational level for hydrophilic broadleaf species.

### Why is the safety-efficiency trade-off not relevant for all species?

Our data clearly demonstrate that the safety-efficiency trade-off does not affect individuals of all species equally. Rather, we found considerable differences between the three studied species groups, which credibly differed in all three analysed Tichý-Ellenberg indicator values, indicating pronounced differences in habitat preference.

As hypothesized, in conifers we only found a within-species relationship between safety and efficiency. In gymnosperms, the tracheid-based wood anatomy limits the maximum attainable *K*_s_ (Becker et al., 1999; Tyree & Zimmermann, 2002). Although gymnosperms use a multitude of strategies to mitigate the effect of smaller conduits, such as the torus-margo pit morphology that permits higher flow rates without compromising safety (Pittermann et al., 2005), higher conduit densities, and adjustments to the leaf-to-sapwood-ratio (Becker et al., 1999), they on average achieve a lower maximum conductivity per sapwood area than angiosperms. The conduit anatomy of conifers renders higher values of *K*_s_ unavailable to selection and hence poses an absolute constraint *sensu* Roff & Fairbairn (2007) on hydraulic efficiency. *P*_50_, on the other hand, is not constrained in that way and in our dataset varied more or less over the same range as in angiosperms (Fig. 5), highlighting the importance of quantities other than conduit diameter in determining embolism resistance (Lens et al., 2022). The restricted range in *K*_s_ results in a lower power to detect existing across-species safety-efficiency relationships in conifers (Bland & Altman, 2011; Isasa et al., 2023) and may completely mask them if accompanied by a ‘grade shift’ (Garland, 2014), i.e., differences in trait averages between species that offset the relationship between traits across species. While this may explain the weak across-species relationship in conifers found here and elsewhere (Willson et al., 2008; Larter et al., 2017; Song et al., 2021), it does not preclude finding across-species patterns in larger datasets, which indeed have been reported in numerous studies (e.g., Pittermann et al., 2011; Gleason et al., 2016; Liu et al., 2021).

**Figure 5.**
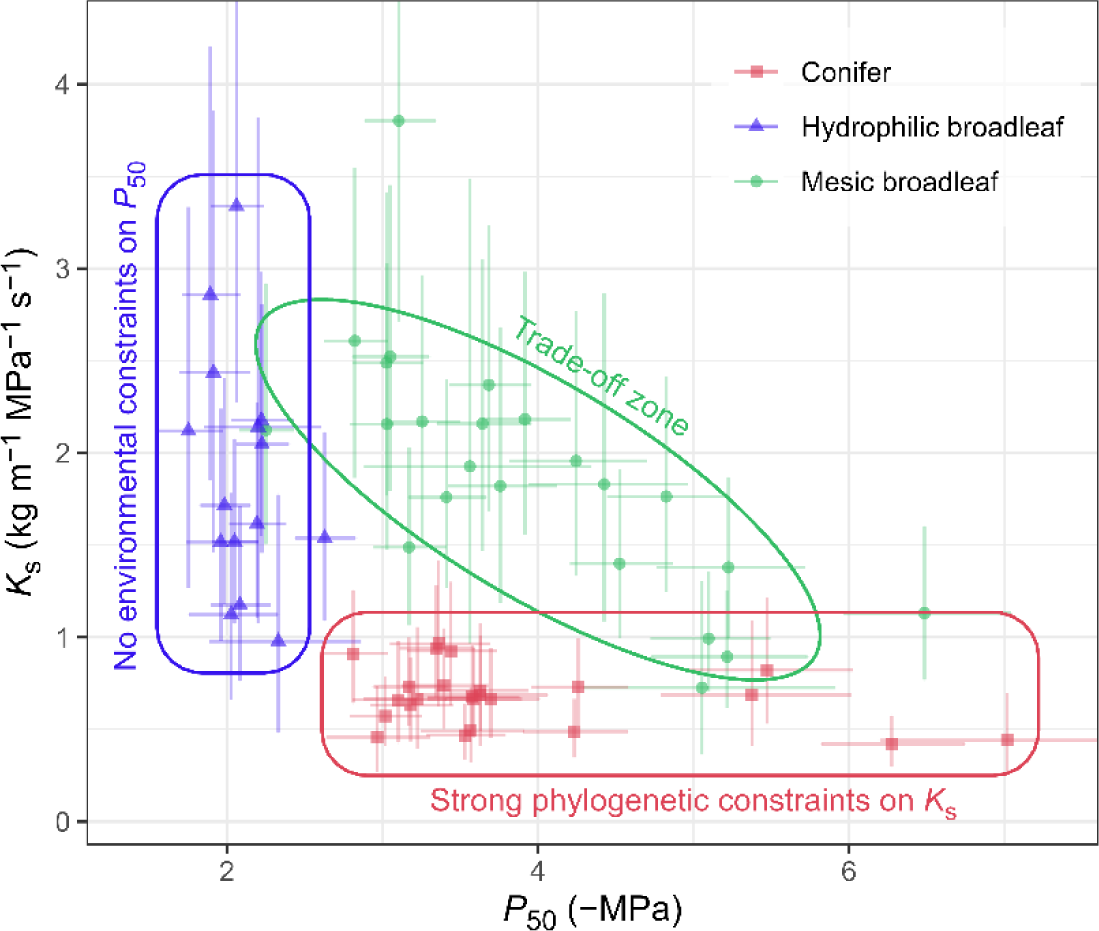
Conceptual visualization of the safety-efficiency trade-off in the studied tree species. Shown are the estimated species means of *P*_50_ and *K*_s_ (posterior means with 95 % credible intervals). Observed data show that most mesic broadleaf trees (green) occupy the theoretical trade-off zone of Gleason et al. (2016), while hydrophilic broadleaves (blue) occupy the range of low embolism resistances, and conifers (red) cluster in a zone of low efficiency.

The observed lack of a safety-efficiency relationship on either organizational scale for hydrophilic species also is in line with our assumptions. Hydrophilic species naturally grow in habitats where water is not a limiting factor, and where adaptations to water-logging and flooding are likely to be far more strongly selected for than adaptations against drought. Moreover, the intense disturbance regime particularly in riparian sites close to the river edge (Bendix et al., 1997; Tabacchi et al., 1998) selects for species with high colonization and regeneration potential, favouring trait combinations characteristic of pioneer species. Species following such a strategy are likely to prioritize fast growth and reproduction under favourable conditions over investments into structures that permit them to withstand unfavourable conditions (Grime, 2002; Laughlin, 2023). For these reasons, there were likely little to no environmental constraints on hydraulic safety in the evolution of the wood anatomy of the hydrophilic species, resulting in a relatively low embolism resistance that is decoupled from *K*_s_ for all species of that group (Fig. 5). This is consistent with the notion that trade-offs are most observable under fixed and limited amounts of resources, while additional resource acquisition may mask existing trade-offs (van Noordwijk & de Jong, 1986; Agrawal, 2020). In agreement, Liu et al. (2021) found that low seasonality and wet habitats may push species towards trait combinations with low efficiency and low safety.

The only group where we found evidence for both within-and across-species relationships between *P*_50_ and *K*_s_ were mesic broadleaved species. While this group bundles species from many different habitats, they are united by the fact that they neither grow in habitats with a weak selection for embolism safety nor are evolutionarily restricted to a reduced range in conductive efficiency. Hence, they can be expected to be fully affected by the safety-efficiency trade-off, placing them squarely in the “trade-off zone” in Fig. 5. The observed differences in the relationship between the three species groups clearly demonstrate that to understand how a trade-off affects individuals of different species and why some are pushed away from optimal trait values towards ‘incompetent’ trait combinations, both environmental and evolutionary constraints have to be taken into account.

### How does the observational scale affect safety-efficiency relationships?

The observable patterns in *P*_50_-*K*_s_-relationships strongly depend on the observational scale. The mechanisms commonly assumed to be responsible for the safety-efficiency trade-off act on the pit membrane, conduit and/or tissue level. The presence of a functional or allocation constraint on those smaller scales does not necessarily result in an observable negative trait association on the individual level, as plants possess considerable ‘degrees of freedom’ (Garland, 2014) in their wood anatomy that can offset such a relationship (Meinzer et al., 2010; Grossiord et al., 2020), e.g. by compensating reductions in hydraulic efficiency due to an increased pit membrane thickness by modifications in vessel density or connectivity that affect embolism safety in a different manner. In consequence, even within species, the existence of a trade-off on pit membrane or conduit level does not always imply a negative phenotypic correlation on individual level. In addition, it is not guaranteed that patterns resulting from processes acting on the individual level will also be visible on the species level, as species-specific adjustments of the aforementioned degrees of freedom may mask existing within-species patterns. For the same reason, the negative correlation between safety and efficiency may be stronger between phylogenetically more closely related species that share a similar wood anatomy (Gleason et al., 2016).

An important consequence of the fact that selection acts on the individual level is that observed trait correlations at higher organizational scales cannot be interpreted in the same way as within-species patterns. While trait correlations based on species means can be interpreted as the *outcome* of evolutionary processes, they neither reflect current constraints nor future evolutionary trajectories (Futuyma & Moreno, 1988; Armbruster & Schwaegerle, 1996; Agrawal, 2020), complicating the attribution of observed negative trait correlations to specific mechanisms. In addition to the constraints acting on individuals, across populations or species, trait associations are affected by processes that can result in higher-level patterns that differ from within-population patterns, such as correlational selection (Armbruster & Schwaegerle, 1996; Roff & Fairbairn, 2012) or niche specialization (Agrawal, 2020). Moreover, selection can also break associations between traits that are present at lower levels, e.g., when speciation results in fundamental changes in strategies (Schluter, 1996; Agrawal, 2020). Accordingly, species-level correlations between safety and efficiency could be driven by the evolutionary legacy of the functional and allocation constraints on pit membrane, conduit and tissue level commonly thought to be responsible for the trade-off, but as well result from processes acting at higher scales.

If the safety-efficiency trade-off was driven by mechanistic constraints on the xylem level, we would expect to find a stronger correlation between safety and efficiency within than across species. Our data did not confirm this for any of the analysed species groups. A potential reason for this is that as in our study there was no replication within individuals, intra-individual variability in *P*_50_ and *K*_s_ was subsumed into the within-species variance, biasing the within-species correlation between the two variables towards zero. Further, the common garden design of the experiment aimed at minimizing the effect of environmental heterogeneity may have resulted in reduced trait ranges, though simultaneously excluding the effect of potentially confounding trait-environment relationships. For the analysed conifers, the within-species correlation was likely further weakened as here *P*_50_ and *K*_s_ had to be measured on different segments. However, it seems probable that the reason for the stronger across-species correlations is that the trade-off is at least in part driven by processes acting at higher scales, particularly niche specialization aligning traits along axes of advantageous trait combinations, resulting in a strategic trade-off *sensu* Agrawal (2020). Similar processes seem to be at play, e.g., for the traits of the worldwide leaf economics spectrum, where quantitative genetic data imply that selection played a larger role in shaping across-species patterns than hard genetic constraints (Donovan et al., 2011). In the context of the safety-efficiency trade-off, the constraints at the xylem level may merely define a boundary line (Grubb, 2016) that sets the bounds for *P*_50_ and *K*_s_ in a limit relationship (cf. Bittencourt et al., 2016), with lower values of *P*_50_ and *K*_s_ being available to selection. ‘Incompetent species’ (Liu et al., 2021) in that context would arise from selection, resulting from a coordination of safety and efficiency with other traits that confer advantages at sub-optimal values of *P*_50_ and *K*_s_ in a whole-plant water-use strategy. In any case, the context-dependent character of the safety-efficiency relationship and its inconsistency across scales found in this study suggests that environment-driven selection plays a substantial role for the observed across-species patterns.

### Why is the evidence about the safety-efficiency trade-off so inconsistent?

While the observed strength of the across-species correlation of the pooled dataset (R^2^ = 0.18) is in line with values found in the literature (angiosperms: R^2^ = 0.10; gymnosperms: R^2^ = 0.15; Liu et al., 2021), the across-species correlation for the mesic broadleaved species (R^2^ = 0.49) is substantially higher. While Gleason et al. (2016) reported similarly strong correlations within certain families, such a high correlation across species from different families is remarkable. The main reason for the strong across-species pattern in these species compared to other studies is likely the exclusion of species that are not affected by the trade-off in the same way due to constraints resulting from their evolutionary history or habitat preferences (cf. Fig. 5). However, our study differs from the larger meta-studies also in the fact that all species were sampled following a consistent protocol on evenly aged trees grown together in a common garden and measured in the same laboratory with the same methods. By using consistent protocols, we bypassed comparability issues between different vulnerability curve methods, particularly since we omitted ring-porous species which are affected by vessel-length related measurement artefacts (Cochard et al., 2010; Torres-Ruiz et al., 2014; Schuldt et al., 2026). Furthermore, we used consistent methods for estimating *K*_s_, where values may differ by a factor of two just depending on the selection of the cross-sectional area used for normalization (cf. Hoeber et al., 2014; Hajek et al., 2014). The controlled design in our dataset further minimizes the effects of environmental heterogeneity and age-specific patterns on the traits. Notwithstanding these methodological advantages, we observed relatively weak average within-species correlations for mesic broadleaved trees and conifers (R^2^ = 0.11 and 0.12, respectively), which suggests that hard constraints on the xylem level may play a smaller role for safety-efficiency relationship than environmentally driven selection. As a large fraction of the existing literature focuses on across-species patterns, which appear to be strongly context dependent, and the existing work on within-species patterns often focuses on relatively small numbers of species, it is not surprising that authors tend to come to vastly different conclusions regarding the importance of the safety-efficiency trade-off.

### Where to go from here?

The multi-scale approach via multi-response mixed models proposed in this paper can be extended in various ways to improve inference on the safety-efficiency trade-off. Firstly, it has been hypothesized that safety and efficiency do not trade off exclusively with each other, but are also involved in trade-offs with other traits, particularly mechanical strength and storage (Baas et al., 2004; Pratt & Jacobsen, 2017). This can result in trait combinations that are pushed away from the theoretical ideal if selection only acted on *P*_50_ and *K*_s_ (Brodersen, 2016). The relationship between safety and efficiency has even been assumed to be a mere consequence of the trade-off between *K*_s_ and mechanical stability and a synergy between mechanical stress and hydraulic safety rather than resulting from a separate trade-off (Bittencourt et al., 2016). The model proposed in this paper can easily be modified to include additional variables describing mechanical stability and/or storage and their covariation with safety and efficiency. This can also be extended into designs that move away from the exclusive study of trade-offs in woody tissues onto the coordination between multiple hydraulic traits across organs in an integrated, whole-plant approach (Meinzer et al., 2010; Brodersen, 2016).

Further, to obtain stronger evidence on the stability-efficiency trade-off within species, more principled tests for genetic correlations between *P*_50_ and *K*_s_ would be advisable. By adopting an appropriate design using replicate measurements in common garden experiments across different sites, the phenotypic and genetic components of the correlation between traits can be separated. An example of a study focusing on phenotypic plasticity in the safety-efficiency trade-off can be found in Pritzkow et al. (2020), though these authors neither did find evidence for a trade-off nor did they compute genetic correlations. Given an appropriate design, our model can be extended to allow a decomposition of phenotypic covariance into heritable and not heritable components (Dingemanse & Dochtermann, 2013; Laitinen & Nikoloski, 2024).

Finally, moving onto larger evolutionary scales, it would be important to study to which degree the trade-off between safety and efficiency is phylogenetically conserved. An analysis of this type (Sanchez-Martinez et al., 2020) on the genus and species level found no evidence for a phylogenetically conserved correlation between safety and efficiency. Again, it is straightforward to extend our model to a multi-response phylogenetic mixed modelling framework (Westoby et al., 2023; Halliwell et al., 2025) to combine both tests of coordinated evolution across species and tests of intraspecific residual correlations.

### Conclusions

Our results demonstrate that the xylem safety–efficiency trade-off is not universal but strongly depends on ecological context, evolutionary constraints, and observational scale. We were able to confirm our hypotheses that safety and efficiency in temperate trees are negatively correlated both within and across species, but that this correlation is mostly driven by mesic broad-leaved species, which are neither affected by the evolutionary constraints on *K*_s_ faced by conifers nor released from selective pressure on embolism resistance like hydrophilic broad-leaved species. However, contrary to our expectation, within-species correlations were weaker than across-species correlations, which would be expected if functional or allocation constraints on the pit membrane, conduit or tissue level were the main driver of the trade-off. Instead, our findings indicate that the broader cross-species patterns at least partially result from ecological differentiation and niche specialization among species rather than from mechanistic constraints on the xylem level. The differences across scales pinpoint the relevance of considering the observational scale for the analysis of evolutionary trade-offs, as the same data can lead to opposing conclusions depending on which level of organization they are analysed on. Ultimately, to obtain more reliable inference on the safety-efficiency trade-off, future studies will have to combine data from several data sources using species spanning different habitats and ecological strategies, and analyse them on multiple scales of aggregation while accounting for their ecological and evolutionary context.

## Acknowledgements

We wish to express our gratitude for the support of Tom Künne during sample collection in the ARBOfun arboretum. RML further wishes to thank Kerstin Pierick for valuable discussions and BS&G for moral support.

## Author contributions

BS developed the original research questions, which were modified by RFT and RML to the fit the scope of the present manuscript. RFT performed all physiological measurements and processed the vulnerability curve data. RFT and RML performed the statistical analysis and wrote a first draft, which was then intensively discussed and revised by the all authors. RFT and RML share equal first authorship of this work.

## Data availability statement

The species level dataset is available in Supplementary Table S1. Individual level data can be made available upon request.

### Boxes

**Box 1: Mechanistic explanations for the safety-efficiency trade-off**

A number of mechanisms to explain the safety-efficiency trade-off have been proposed that act on different levels of organization, ranging from the level of individual pit membranes over the level of single conduits to the tissue level.

#### Pit membrane level

On the level of individual pit membranes between conduits, a functional trade-off between safety and efficiency is likely, as both hydraulic conductance (Kaack et al., 2019) and the spread of emboli depend on the same pit properties, namely pit membrane thickness, area and porosity. This notably is true both for concepts of embolism spread driven by gas diffusion (Pereira et al., 2023) and for classical air seeding concepts via bulk flow of gas bubbles (Christman et al., 2009). For that reason, on the pit membrane level, any increase in safety is likely to result in a concomitant reduction in efficiency.

#### Conduit level

On the level of single conduits, a potential safety-efficiency trade-off has often been hypothesized, though its existence largely depends on the assumption that embolism risk is related to conduit size. While such a link is well established for frost-induced embolism (Ewers, 1985; Hacke & Sperry, 2001; Mayr & Sperry, 2010), its existence remains debated for embolism resulting from drought (Anfodillo & Olson, 2021; Lens et al., 2022; Isasa et al., 2023). The classical mechanistic explanation for a conduit-level trade-off is based on the pit-area hypothesis, which posits that while longer and wider conduits achieve a higher hydraulic conductance, they also have a higher total area of interconduit pits and hence are more likely to possess a pore large enough to allow embolism formation via air-seeding (Wheeler et al., 2005; Hacke et al., 2006; Christman et al., 2009). However, leaky pit membranes with pores large enough to permit substantial mass flow of gas are likely not common enough to drive embolism spread in the way predicted by this hypothesis (Kaack et al., 2021). Instead, embolism spread may be predominantly driven by gas transport via diffusion (Guan et al., 2021) or in the form of stable nanobubbles (Jansen et al., 2026), which is not constrained to rare leaky pits and therefore not obviously affected by conduit dimensions. However, recent modelling by Pereira et al. (2023) suggests that diffusion-driven embolism formation may also depend on conduit size. Provided a link between conduit size and embolism risk exists, a higher efficiency due to increased conduit diameters will result in a reduction in safety.

#### Tissue level

On larger spatial scales, the link between safety and efficiency on pit membrane and conduit level is compounded by the effect of the connectivity of the conduit network formed by the entirety of conduits connected by interconduit pits. A higher connectivity has been shown to permit higher flow rates (Loepfe et al., 2007; Martínez-Vilalta et al., 2012). At the same time, higher network connectivity facilitates the spread of embolism to adjacent conduits (Loepfe et al., 2007; Johnson et al., 2020; Mrad et al., 2021). In addition, on the tissue level, conductance can be increased by increasing the fraction of area occupied by conductive tissue, which can result in higher embolism risk due to a lower allocation into tissues able to withstand mechanical stresses under low water potentials (Bittencourt et al., 2016), or due to faster diffusive gas transport to close-by conduits (Guan et al., 2021). Hence, there are several processes on tissue level that could result in a negative safety-efficiency relationship.

## Supporting information

### Notes S1. Supplementary methods

The following document describes the full structure of the multi-response mixed model for the trade-off between embolism safety (−*P*_50_) and hydraulic efficiency (*K*_s_).

The model was fitted in `R` v. 4.5.2 (R Core Team, 2025) using `R` package `brms` v.2.23.0 (Bürkner 2017; 2018).

### S1.1 Structure of the trade-off model

To analyse the safety-efficiency trade-off within and across species, we modelled natural log-transformed negative *P*_50_ (*_x_*_[*ij*]_) and natural log-transformed *K*_s_ (*y*_[*ij*]_) for observation *i* of *I* belonging to species *j* of *J* jointly in a multi-response mixed model. First, the estimation uncertainty in the *P*_50_ estimates from Eqn. (1) was included explicitly in our model using a measurement error model for *x*_[*ij*]_:

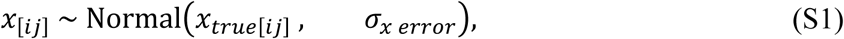

where *x_true_*_[*ij*]_ is the unmeasured true value of *x*_[*ij*]_ and *σ_error_* is the estimation uncertainty in log(*P*_50_). As *P*_50_ is estimated in the model in Eqn. (1) on the untransformed scale, the standard error *σ_x error_* on the transformed was estimated from the standard errors *SE* and estimates *Est* on the untransformed scale as *σ_x error_* = √log((*SE*/*Est*)^2^ + 1). We then jointly modelled *x_true_*_[*ij*]_ and *y*_[*ij*]_ as:

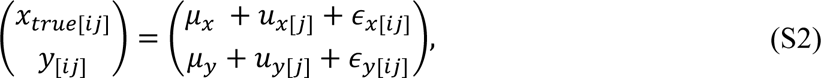

where *μ_x_* and *μ_y_* are the grand means, *u_x_*_[*j*]_ and *u_y_*_[*j*]_ are species-level random intercepts and *ε_x_*_[*ij*]_ and *ε_y_*_[*ij*]_ are the individual-level residuals for *x_true_*_[*ij*]_ and *y*_[*ij*]_, respectively.

The species level random effects were assumed to follow a multivariate normal distribution with mean 0 and covariance matrix Σ*_across_*:

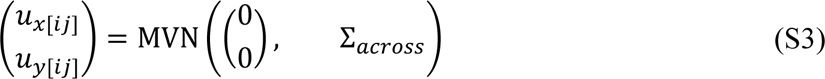

Analogously, the individual-level residuals were drawn from a multivariate normal distribution with mean 0 and covariance matrix Σ*_wit_*_ℎ*in*_:

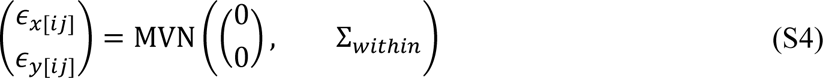

The covariance matrix Σ_across_ can be decomposed into two 2 × 2 diagonal matrices with the standard deviations *τ_x_* and *τ_y_* and a 2 × 2 correlation matrix Ω_across_ containing the across-species correlation *ρ_across_*:

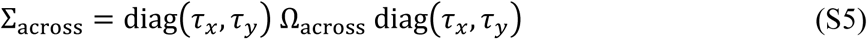

Analogously, the covariance matrix Σ_across_ can be decomposed into two 2 × 2 diagonal matrices with the standard deviations *σ_x_* and *σ_y_* and a 2 × 2 correlation matrix Ω_within_ containing the within-species correlation *ρ_wit_*_ℎ*in*_:

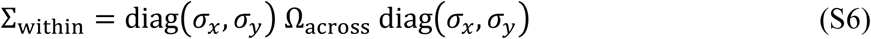

### S1.2 Choice of priors

To simplify prior choices, these were expressed on the scale of the data, using the sample mean (*M_x_* and *M_y_*) and standard deviation (*SD_x_* and *SD_y_*) of for *x*_[*ij*]_ and *y*_[*ij*]_, respectively. The grand means *μ_x_* and *μ_y_* were assigned a normal priors centred around the mean of the response variable with a standard deviation equal to half the standard deviation of the response variable:

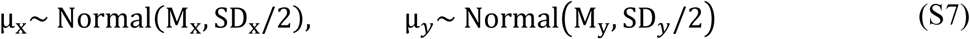

This prior constrains the grand means to the range of the data, slightly pulling them to the centre.

The standard deviations for the species-specific random effects *τ_x_* and *τ_y_* were assigned half-normal priors with a standard deviation equal to the standard deviation of the response:

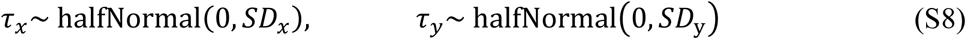

The residual standard deviations *τ_x_* and *τ_y_* analogously were assigned half-normal priors:

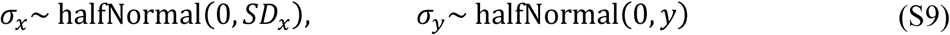

Finally, the correlation matrices Ω_across_ and Ω*_wit_*_ℎ*in*_ were assigned *LKJ*(2) prior (Lewandowski, Kurowicka & Joe, 2009), which slightly pushes the correlations away from the boundaries while not ruling out any other potential values:

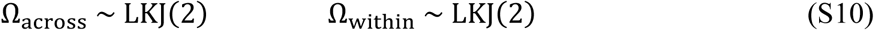

The same prior choices (except for the scales of the data) were used for the model pooling across all species and the three models for separate species groups.

### S1.3 Model fitting and evaluation

All models were fitted on 4 parallel chains with 10000 iterations each, the first 5000 of which were discarded as warm-up. For all models, we used a target acceptance rate of `adapt_delta = 0.9` and a maximum tree depth of 12. Additionally, we constrained starting values to ±0.5 on the unconstrained scale by setting `init_r = 0.5` and reduced the initial stepsize to 0.1.

All model parameters of all models reached an *R̂*-value of below 1.002832 and a total effective sample size of at least 5902, indicating full model convergence and sufficiently large posterior samples for all parameters. As there was a small fraction of observations that reached pareto k values of above 0.7 in approximate leave-one-out cross-validation based on Pareto-smoothed importance sampling with the loo package (Vehtari et al., 2017, 2025), loo-based leave-one-out cross-validation was performed with re_loo = TRUE to guarantee reliable estimates of the expected log pointwise predictive density.

### S1.4 Derived quantities

Based on the parameter estimates of our model, we calculated the standardized major axis (SMA) regression slopes and intercepts for the relationship *P*_50_ and *K*_s_ across and within species. The SMA slope is equal to the sign of the correlation between two variables multiplied with the ratio of their standard deviations (Warton et al., 2006). We thus calculated the slope *β_across_* of the across-species relationship as:

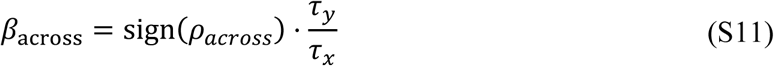

The slope *β_wit_*_ℎ*in*_ of the within-species relationship can be calculated in the same form as:

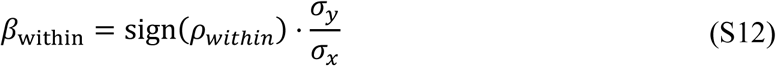

The intercept *α_across_* of the across-species relationship can be calculated as the mean of *y* minus the mean of *x* times the slope (Warton et al., 2006):

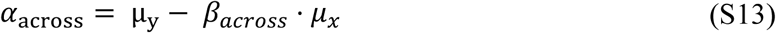

The intercept *α_wit_*_ℎ*in*[*j*]_ of the across-species relationship is species-specific and can be calculated for each species *j* in *J* as follows:

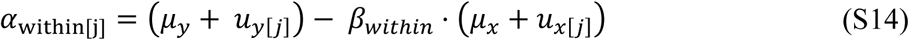

As the SMA slope depends on the sign of the correlation, its posterior distribution becomes bimodal if the posterior distribution of the correlation coefficient does not exclude zero. For that reason, the credible intervals for the SMA slopes and SMA predictions have to be taken with care in these cases, and were excluded (Fig. 3) or highlighted (Fig. S4) in the corresponding figures.

For the expected values of *P*_50_ and *K*_s_ on the untransformed scale (Table S4, S5), we corrected for retransformation bias (Smith, 1993) resulting from Jensen’s inequality by adding half the across-species and residual variance before exponentiating:

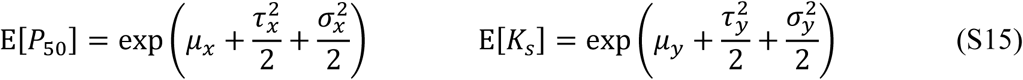

To provide a Bayesian equivalent to the frequentist p-value, we computed the probability of direction *pd* according to Makowski et al. (2019) for all relevant estimates of parameters that are not bound at zero. The probability of direction *pd* is the proportion of the posterior distribution that has the same sign as the median of the posterior distribution.

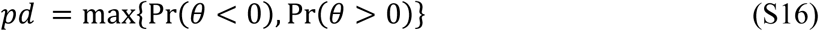

We then calculated a two-sided Bayesian pseudo-p-value *P_Bayes_* as follows:

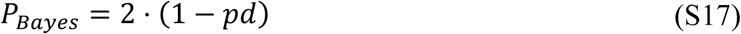

### S1.5 Variance decomposition and total correlation

The variance in *x_true_* and *y* can be decomposed into the variance explained by across-species and within-species differences in the traits:

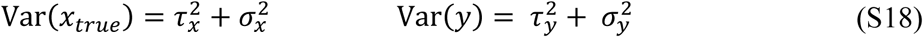

As the fraction of shared variance between correlated variables is the square of their correlation coefficient, this variance can be further decomposed into components that are shared between*x_true_* and *y* and components that are unique to each of the two variables:

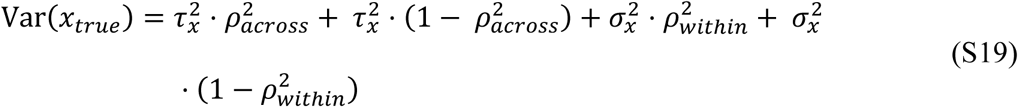

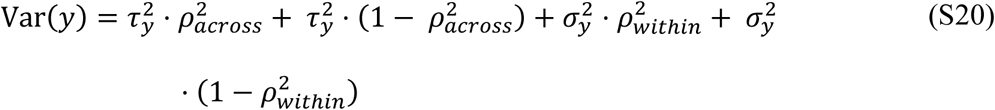

The total covariance between *x_true_* and *y* is equal to the sum of the covariances within and across species:

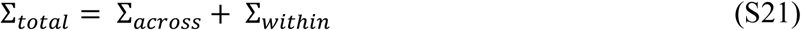

This can be used to compute the total correlation *ρ_total_* between *x_true_* and *y* by dividing the off-diagonal elements in Σ*_total_* by the square root of the product of the diagonal elements:

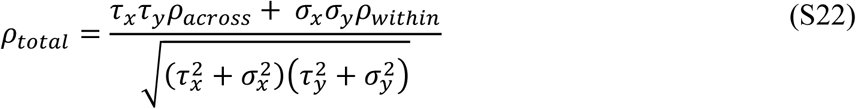

### S1.6 Implementation in brms

The model in S1.1 can be implemented in brms by using the following formula structure:

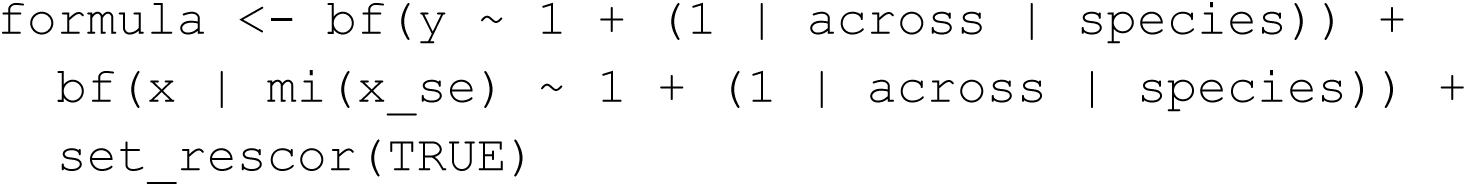

Here, `x` and `y` are the natural log-transformed response variables and `x_se` is the standard error of the estimates of `x`. The `mi()` notation allows specifying for measurement errors and missing data. Setting the species-level random effects with the `across` identifier (or any other label that is consistent between random effects) ensures that a correlation will be estimated between the two random effects vectors. Analogously, `set_rescor(TRUE)` ensures that a residual correlation will be fitted.

This formula can be embedded in a brm() call as follows:

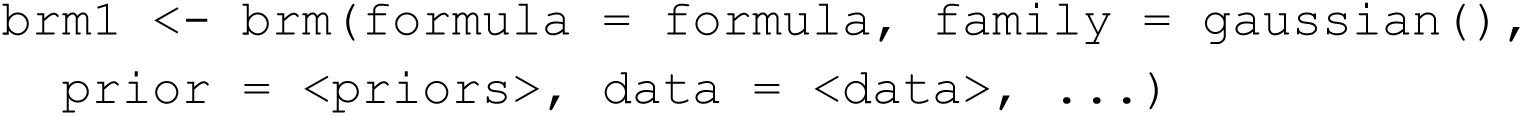

Here, <priors> is a placeholder for an appropriate specification of priors and <data> is a placeholder for the dataset used for modelling.

While it would be appealing to fit a single model with separate parameters by species group, to our knowledge so far this is not possible in brms as residual correlations fitted via set_rescor()cannot be split into groups with different correlation coefficients.

## Supplementary figures

**Figure S1.**
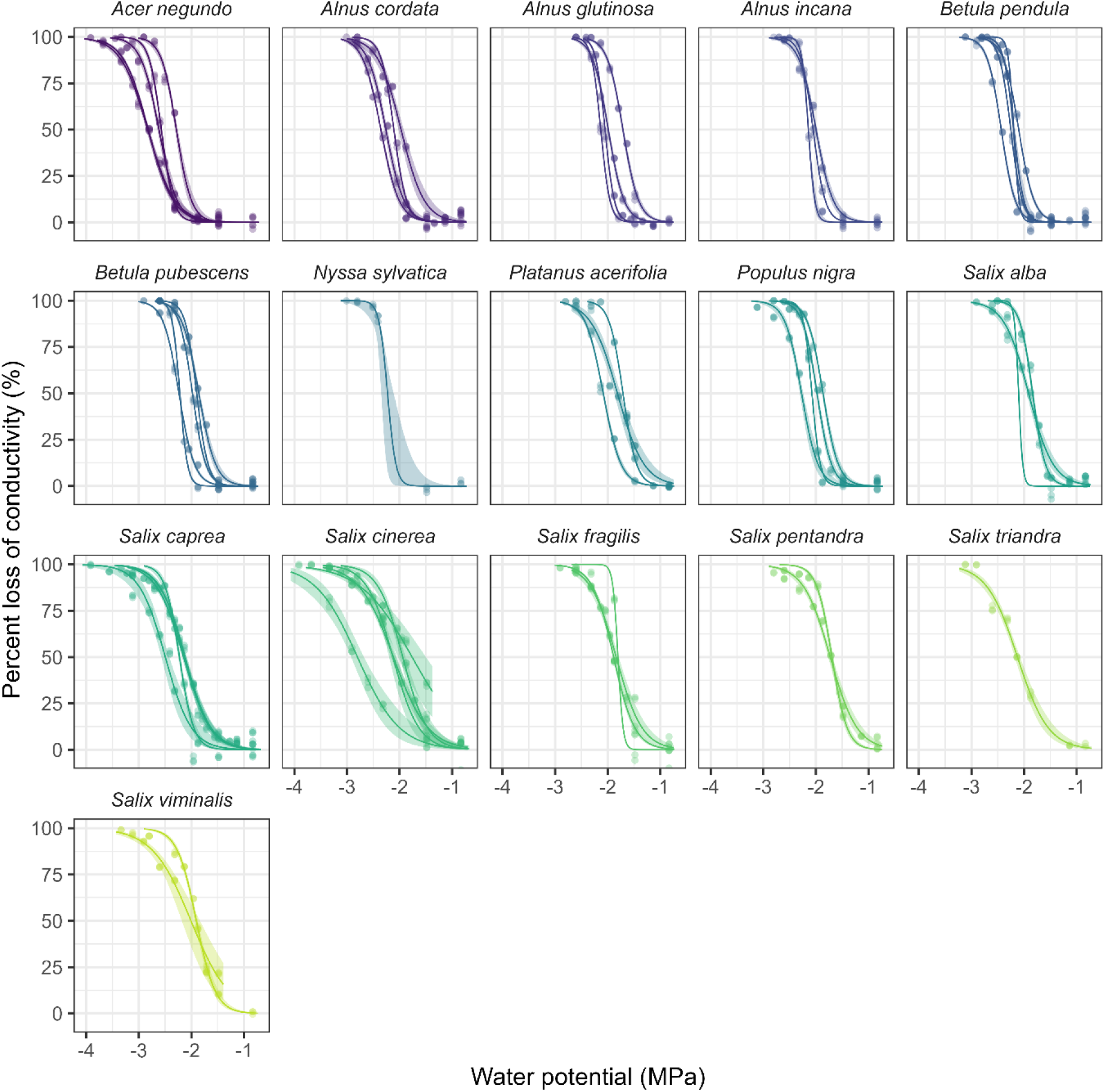
Vulnerability curves for the hydrophilic species. Shown are the observed values (rescaled to percent loss of conductivity scale) overlaid with the predictions (lines) with their 95 % confidence intervals (shaded areas).

**Figure S2.**
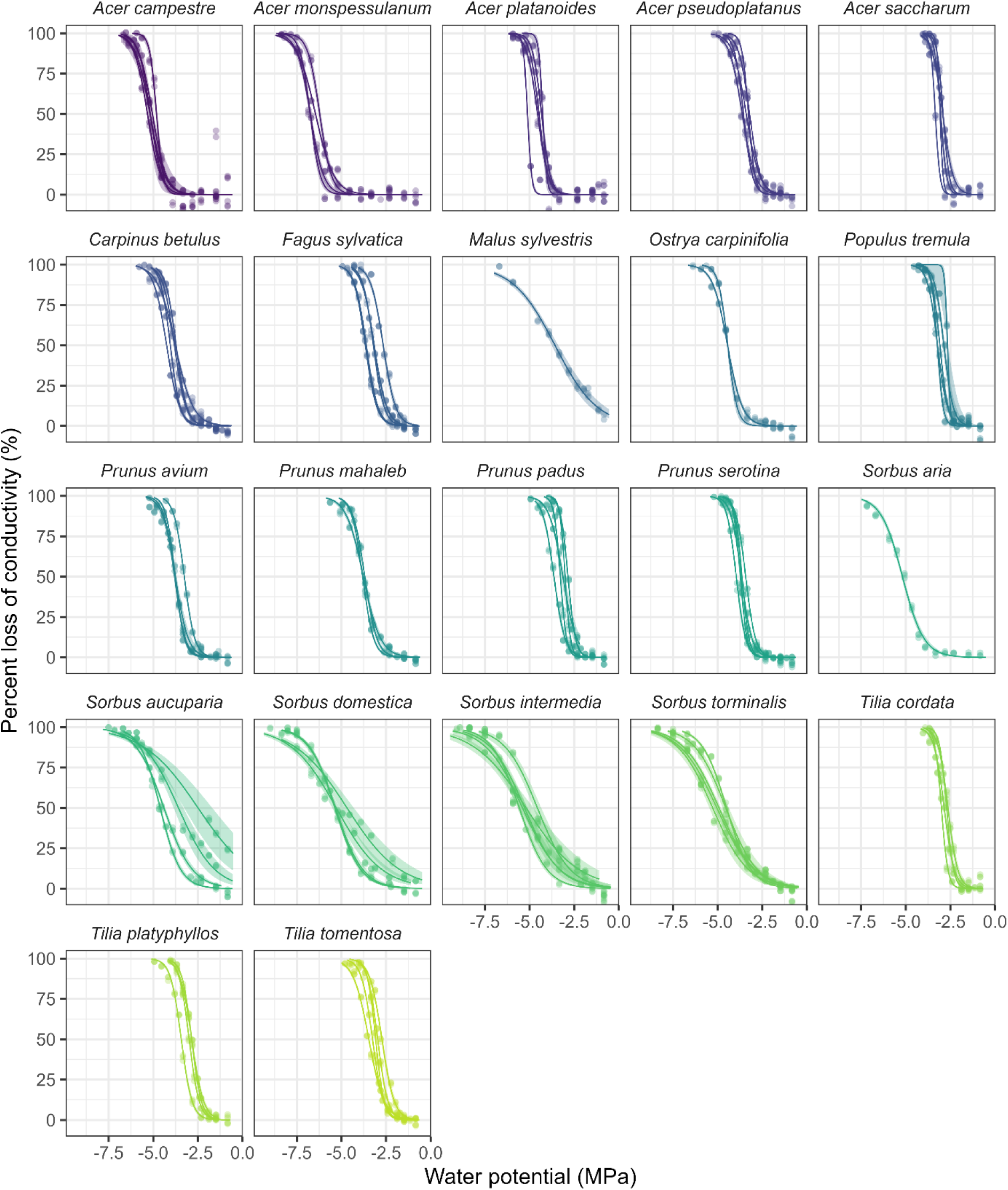
Vulnerability curves for the mesic broadleaf species. Shown are the observed values (rescaled to percent loss of conductivity scale) overlaid with the predictions (lines) with their 95 % confidence intervals (shaded areas).

**Figure S3.**
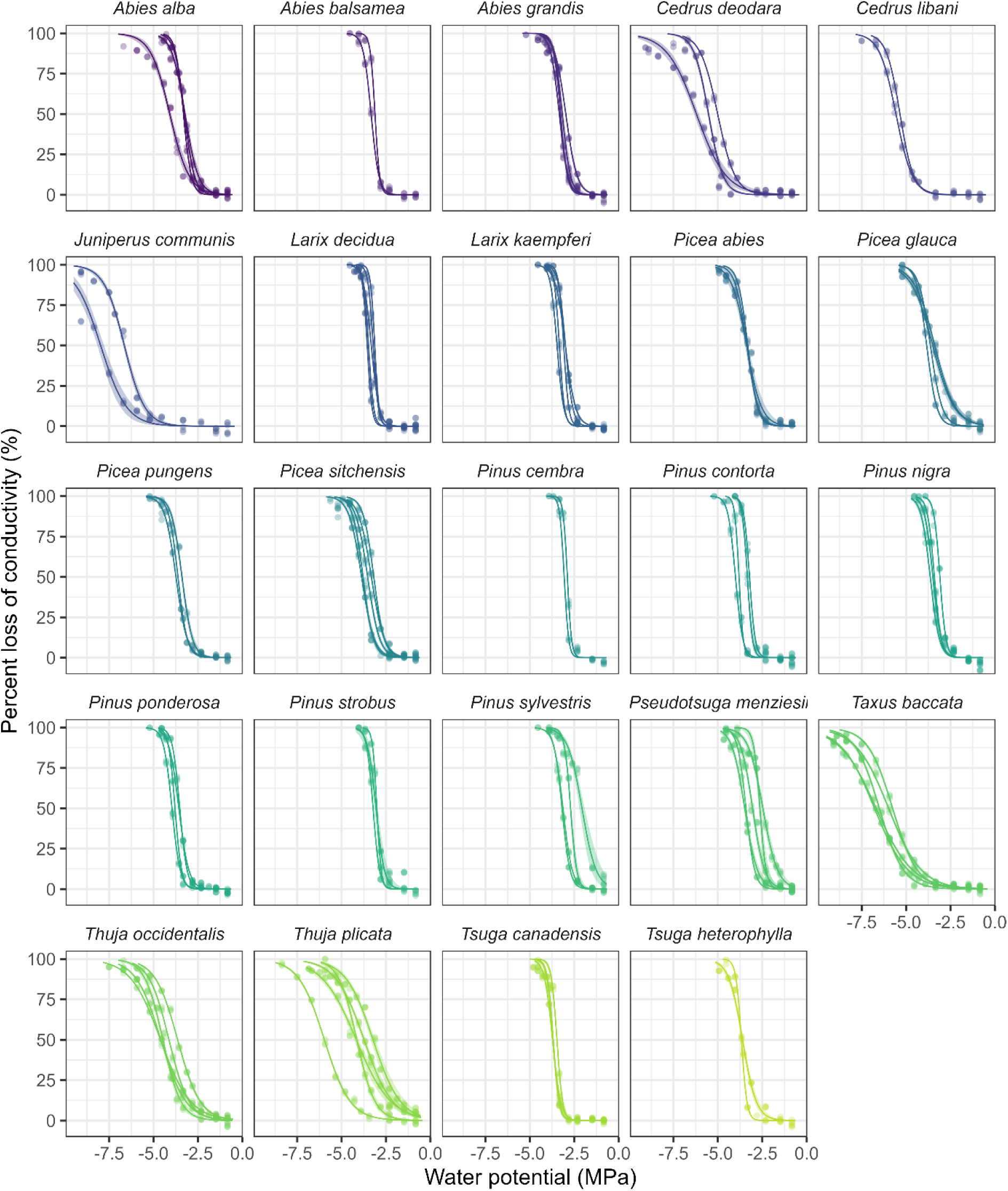
Vulnerability curves for the conifer species. Shown are the observed values (rescaled to percent loss of conductivity scale) overlaid with the predictions (lines) with their 95 % confidence intervals (shaded areas).

**Figure S4.**
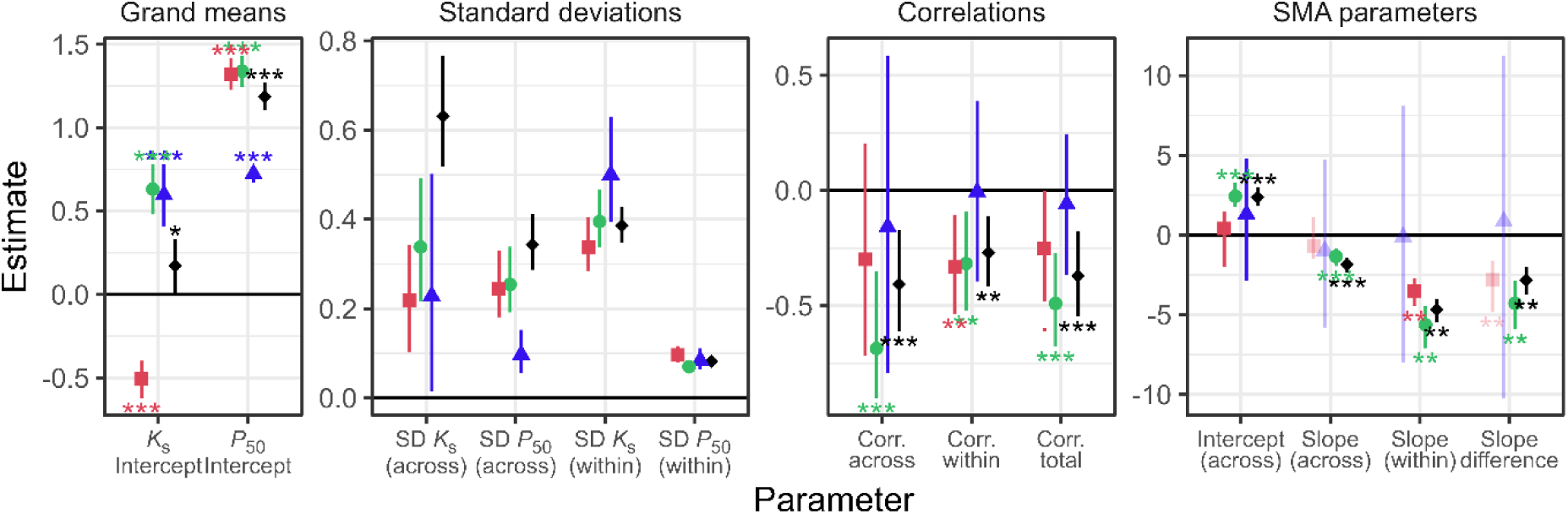
Parameter estimates of the models of the safety-efficiency trade-off. Shown are the posterior mean ± 95 % credible intervals for the model with pooled species groups and the three models with separate parameters per species group groups. From left to right, the four panels show the grand means, standard deviations, correlation parameters and the estimated SMA slopes and intercepts as well as a test for the difference between the across-and within-species slope. SMA slopes and derived quantities based on correlations not significantly different from zero are shown greyed (cf. Notes S1.4).

## Supplementary tables

**Table S1.**
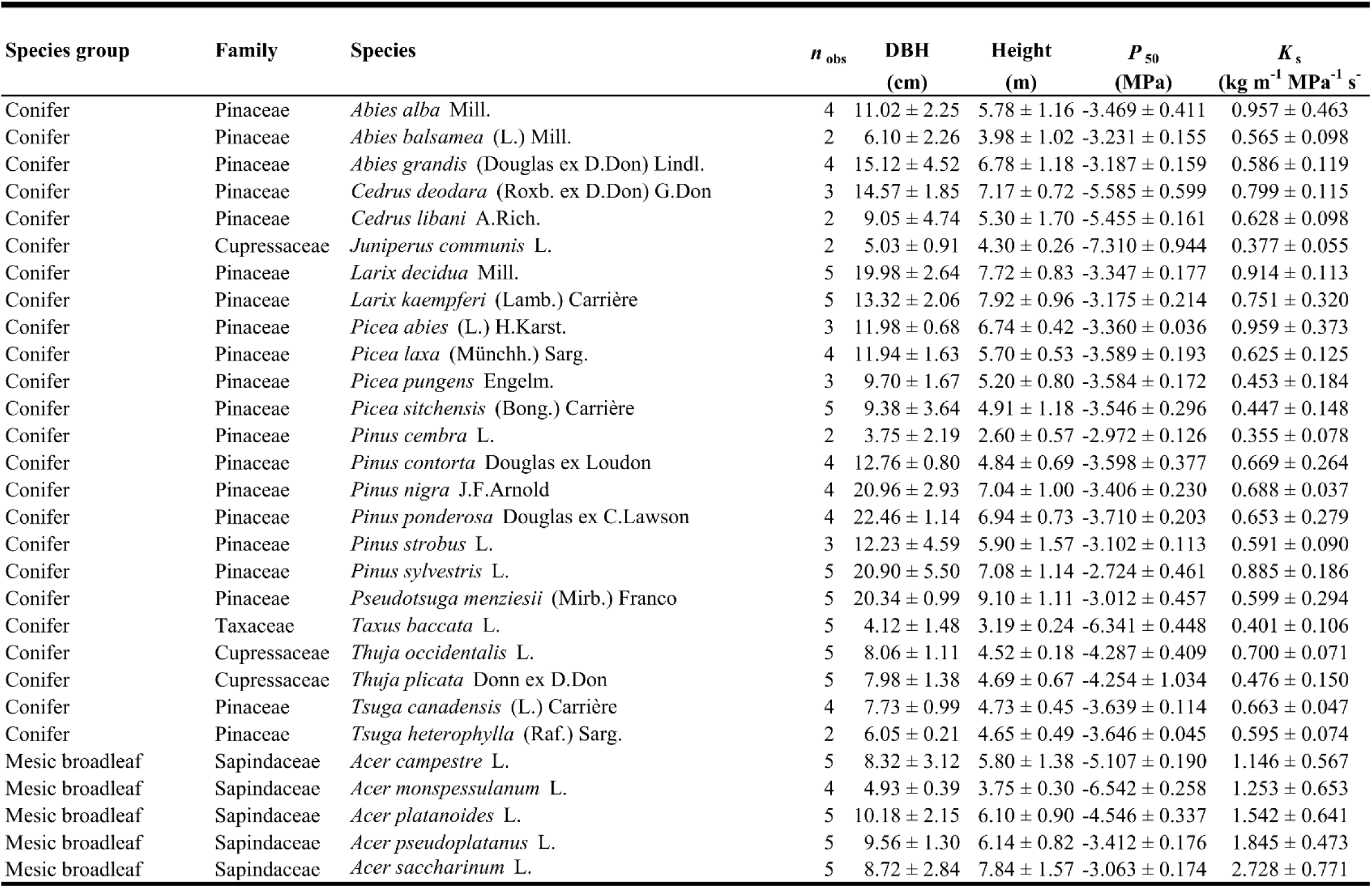
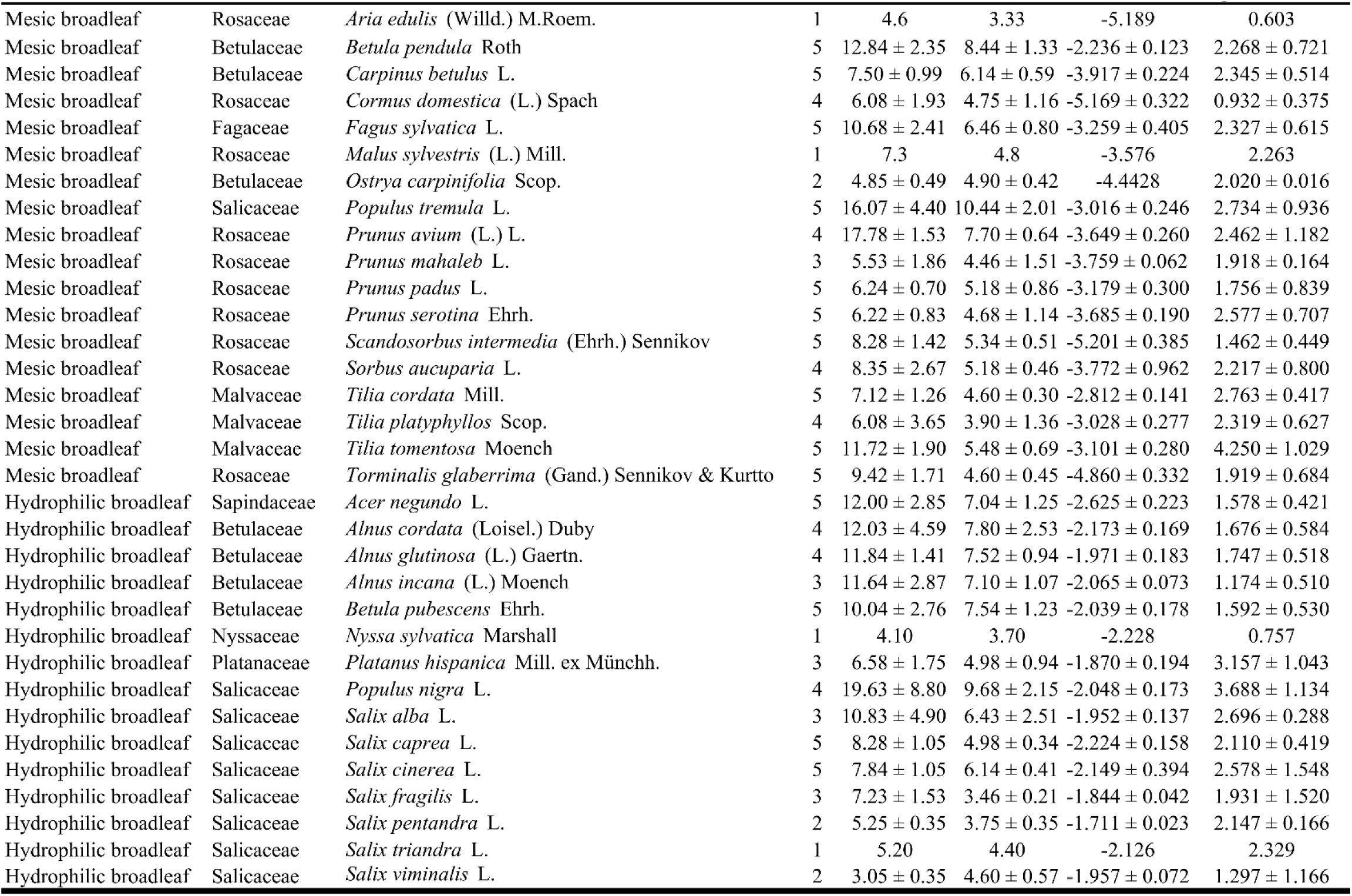
Information about the studied species. Shown are the species group, family, species name, number of and standard deviation of diameter at breast height (DBH), tree height, *P*50 and *K*s.

**Table S2.** Species classified as “hydrophilic” with references for their occurrence and ecological preferences.

| Family | Species | Occurrence |
| --- | --- | --- |
| Sapindaceae | *Acer negundo* L. | Riparian species (Galuszka et al., 2002) |
|  | *Alnus cordata* (Loisel.) Duby | Pioneer species, less dependent on riparian habitats than other *Alnus* species (Pollegioni et al., 2025), diagnostic species in several types of riparian *A. glutinosa* forests (Sciandrello et al. 2022) |
|  | *Alnus glutinosa* (L.) Gaertn. | Riparian forests and swamps (Schnitzler et al., 2007; Stella et al., 2013; Leuschner & Ellenberg 2017) |
|  | *Alnus incana* (L.) Moench | Riparian species (Schnitzler et al. 2007, Hughes et al. 1997; Sburlino et al. 2012; Leuschner & Ellenberg 2017) |
|  | *Betula pubescens* Ehrh. | Swamp forests (Leuschner & Ellenberg; 2017) |
| Nyssaceae | *Nyssa sylvatica* Marshall | Swamp forests (Conner & Day, 1992), floodplain forests (Rheinhardt et al. 1998), tidal freshwater forests (Stahl et al., 2018) |
| Platanaceae | *Platanus* × *acerifolia* (Aiton) Willd. | [horticultural hybrid of the riparian species *P. occidentalis* and *P. orientalis*] |
|  | *P. occidentalis* L. | Riparian forests (Walters & Williams, 1999) |
|  | *P. orientalis* L. | Riparian forests (Stephan & Issa 2017) |
| Salicaceae | *Populus nigra* L. | Riparian forests (Schnitzler et al. 2007, Singer et al. 2013; Leuschner & Ellenberg; 2017) |
|  | *Salix alba* L. | Riparian forests (Schnitzler et al. 2007, Stephan & Issa 2017, Leuschner & Ellenberg; 2017), pioneer species (Leuschner & Ellenberg; 2017) |
|  | *Salix caprea* L. | Pioneer species, in hedges and open sites (Leuschner & Ellenberg; 2017), more mesophile and less tolerant to water-logging than other *Salix* species (Talbot et al., 1987), but can occur in the soft-wood zone of floodplain forest accompanying other *Salix* spp. (Schütt & Stimm; 2004) |
|  | *Salix cinerea* L. | Swamp forests, pioneer species (Leuschner & Ellenberg; 2017) |
|  | *Salix fragilis* L. | Riparian forests, pioneer species (Leuschner & Ellenberg; 2017) |
|  | *Salix pentandra* L. | Swamp forests, pioneer species (Leuschner & Ellenberg; 2017) |
|  | *Salix triandra* L. | Riparian forests, pioneer species (Leuschner & Ellenberg; 2017) |
|  | *Salix viminalis* L. | Riparian forests, pioneer species (Leuschner & Ellenberg; 2017) |

---

**Table S3.**
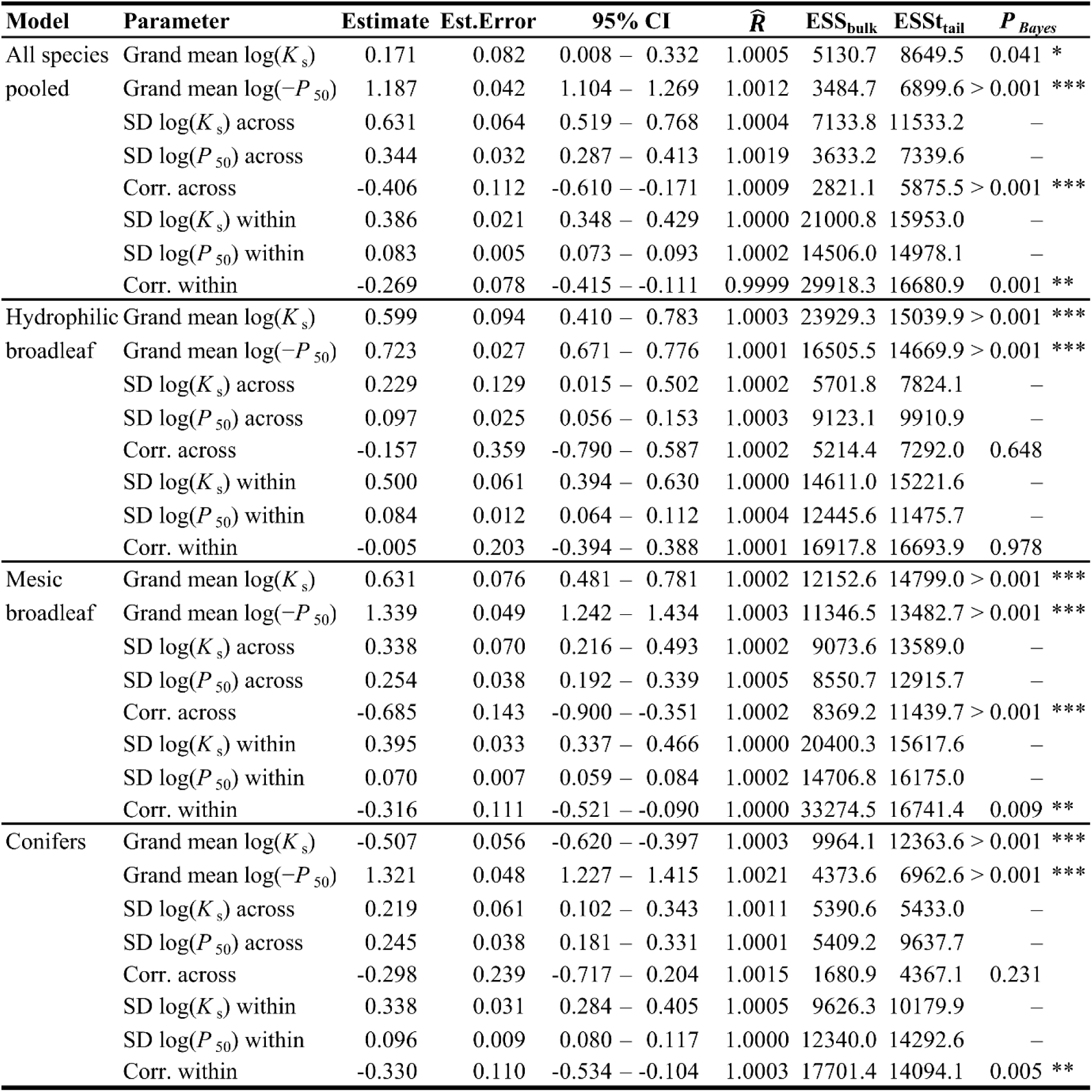
Results of the multi-response mixed models of the relationship between log(−*P*50) and log(*K*s). Shown are the posterior means of the parameter estimates with their MCMC standard error, 95 % credible interval, potential scale reduction factor *R̂*, bulk and tail effective sample sizes and a Bayesian pseudo-p-value *P_Bayes_* (only shown for parameters that are not bounded at zero, see Notes S1.4). Symbols indicate parameters credibly different from zero on the 0.1 (.), 0.05 (*), 0.01 (**) and 0.001 level (***).

| Model | Parameter | Estimate | Est.Error | 95% CI | $\hat{R}$ | ESS<sub>bulk</sub> | ESS<sub>tail</sub> | $P_{Bayes}$ |
| --- | --- | --- | --- | --- | --- | --- | --- | --- |
| All species<br>pooled | Grand mean $\log(K_s)$ | 0.171 | 0.082 | 0.008 – 0.332 | 1.0005 | 5130.7 | 8649.5 | 0.041 * |
| | Grand mean $\log(-P_{50})$ | 1.187 | 0.042 | 1.104 – 1.269 | 1.0012 | 3484.7 | 6899.6 | > 0.001 *** |
| | SD $\log(K_s)$ across | 0.631 | 0.064 | 0.519 – 0.768 | 1.0004 | 7133.8 | 11533.2 | – |
| | SD $\log(P_{50})$ across | 0.344 | 0.032 | 0.287 – 0.413 | 1.0019 | 3633.2 | 7339.6 | – |
|  | Corr. across | -0.406 | 0.112 | -0.610 – -0.171 | 1.0009 | 2821.1 | 5875.5 | > 0.001 *** |
| | SD $\log(K_s)$ within | 0.386 | 0.021 | 0.348 – 0.429 | 1.0000 | 21000.8 | 15953.0 | – |
| | SD $\log(P_{50})$ within | 0.083 | 0.005 | 0.073 – 0.093 | 1.0002 | 14506.0 | 14978.1 | – |
|  | Corr. within | -0.269 | 0.078 | -0.415 – -0.111 | 0.9999 | 29918.3 | 16680.9 | 0.001 ** |
| Hydrophilic<br>broadleaf | Grand mean $\log(K_s)$ | 0.599 | 0.094 | 0.410 – 0.783 | 1.0003 | 23929.3 | 15039.9 | > 0.001 *** |
| | Grand mean $\log(-P_{50})$ | 0.723 | 0.027 | 0.671 – 0.776 | 1.0001 | 16505.5 | 14669.9 | > 0.001 *** |
| | SD $\log(K_s)$ across | 0.229 | 0.129 | 0.015 – 0.502 | 1.0002 | 5701.8 | 7824.1 | – |
| | SD $\log(P_{50})$ across | 0.097 | 0.025 | 0.056 – 0.153 | 1.0003 | 9123.1 | 9910.9 | – |
|  | Corr. across | -0.157 | 0.359 | -0.790 – 0.587 | 1.0002 | 5214.4 | 7292.0 | 0.648 |
| | SD $\log(K_s)$ within | 0.500 | 0.061 | 0.394 – 0.630 | 1.0000 | 14611.0 | 15221.6 | – |
| | SD $\log(P_{50})$ within | 0.084 | 0.012 | 0.064 – 0.112 | 1.0004 | 12445.6 | 11475.7 | – |
|  | Corr. within | -0.005 | 0.203 | -0.394 – 0.388 | 1.0001 | 16917.8 | 16693.9 | 0.978 |
| Mesic<br>broadleaf | Grand mean $\log(K_s)$ | 0.631 | 0.076 | 0.481 – 0.781 | 1.0002 | 12152.6 | 14799.0 | > 0.001 *** |
| | Grand mean $\log(-P_{50})$ | 1.339 | 0.049 | 1.242 – 1.434 | 1.0003 | 11346.5 | 13482.7 | > 0.001 *** |
| | SD $\log(K_s)$ across | 0.338 | 0.070 | 0.216 – 0.493 | 1.0002 | 9073.6 | 13589.0 | – |
| | SD $\log(P_{50})$ across | 0.254 | 0.038 | 0.192 – 0.339 | 1.0005 | 8550.7 | 12915.7 | – |
|  | Corr. across | -0.685 | 0.143 | -0.900 – -0.351 | 1.0002 | 8369.2 | 11439.7 | > 0.001 *** |
| | SD $\log(K_s)$ within | 0.395 | 0.033 | 0.337 – 0.466 | 1.0000 | 20400.3 | 15617.6 | – |
| | SD $\log(P_{50})$ within | 0.070 | 0.007 | 0.059 – 0.084 | 1.0002 | 14706.8 | 16175.0 | – |
|  | Corr. within | -0.316 | 0.111 | -0.521 – -0.090 | 1.0000 | 33274.5 | 16741.4 | 0.009 ** |
| Conifers | Grand mean $\log(K_s)$ | -0.507 | 0.056 | -0.620 – -0.397 | 1.0003 | 9964.1 | 12363.6 | > 0.001 *** |
| | Grand mean $\log(-P_{50})$ | 1.321 | 0.048 | 1.227 – 1.415 | 1.0021 | 4373.6 | 6962.6 | > 0.001 *** |
| | SD $\log(K_s)$ across | 0.219 | 0.061 | 0.102 – 0.343 | 1.0011 | 5390.6 | 5433.0 | – |
| | SD $\log(P_{50})$ across | 0.245 | 0.038 | 0.181 – 0.331 | 1.0001 | 5409.2 | 9637.7 | – |
|  | Corr. across | -0.298 | 0.239 | -0.717 – 0.204 | 1.0015 | 1680.9 | 4367.1 | 0.231 |
| | SD $\log(K_s)$ within | 0.338 | 0.031 | 0.284 – 0.405 | 1.0005 | 9626.3 | 10179.9 | – |
| | SD $\log(P_{50})$ within | 0.096 | 0.009 | 0.080 – 0.117 | 1.0000 | 12340.0 | 14292.6 | – |
|  | Corr. within | -0.330 | 0.110 | -0.534 – -0.104 | 1.0003 | 17701.4 | 14094.1 | 0.005 ** |

**Table S4.**
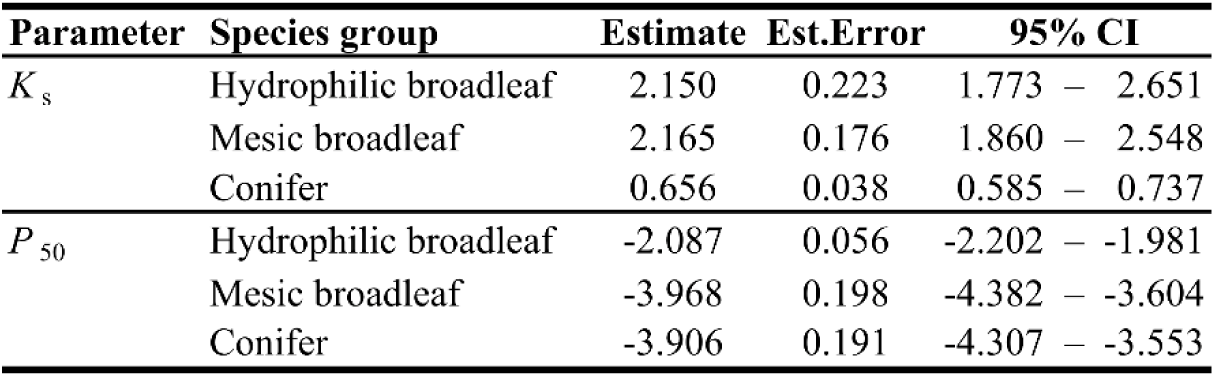
Average estimates of *K*s (kg m^−^¹ MPa^−^¹ s^−^¹) and *P*50 (MPa). Shown are the posterior means of the re-transformed and bias-corrected grand means (cf. Notes S1.4) from Table S2 with their MCMC standard error and 95 % credible interval.

**Table S5.**
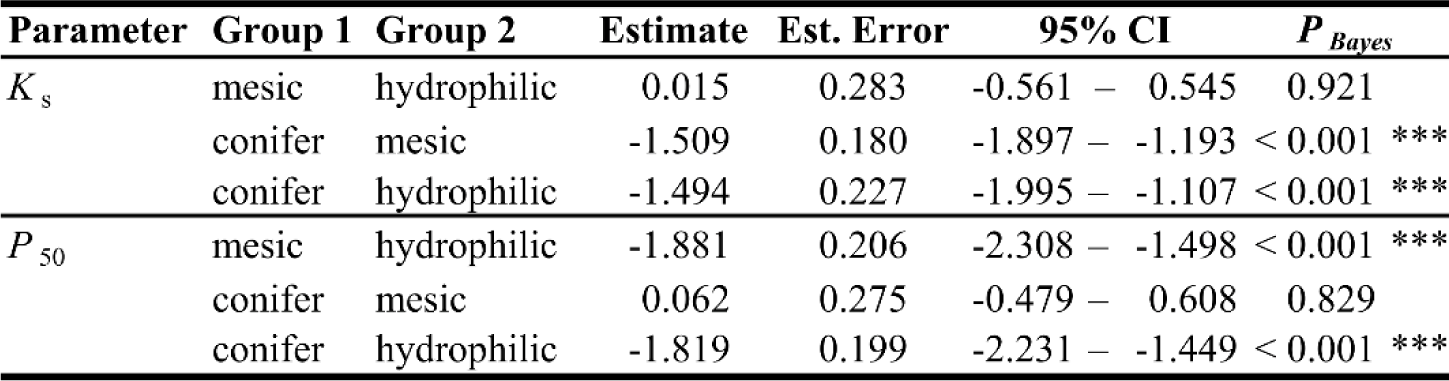
Results of contrast tests of differences the species group means of *K*s (kg m^−^¹ MPa^−^¹ s^−^¹) and *P*50 (MPa). Shown are the posterior means of differences with their MCMC standard error, 95 % credible interval, and a Bayesian pseudo-p-value *P_Bayes_* (see Notes S1.4). Symbols indicate parameters credibly different from zero on the 0.1 (.), 0.05 (*), 0.01 (**) and 0.001 level (***).

**Table S6.** Results of the models of differences in Ellenberg-Tichý indicator values. Shown are the posterior means of the parameter estimates with their MCMC standard error, 95 % credible interval, potential scale reduction factor *R̂*, bulk and tail effective sample sizes and a Bayesian pseudo-p-value *P_Bayes_* (see Notes S1.4). Symbols indicate parameters credibly different from zero on the 0.1 (.), 0.05 (*), 0.01 (**) and 0.001 level (***).

| Model | Class | Parameter | Estimate | Est. error | 95% CI | $\hat{R}$ | ESS<sub>bulk</sub> | ESS<sub>tail</sub> | $P_{Bayes}$ |
| --- | --- | --- | --- | --- | --- | --- | --- | --- | --- |
| Moisture | Fixed | Intercept | 5.117 | 0.311 | 4.508 – 5.734 | 1.0003 | 13763.0 | 12171.2 | < 0.001 *** |
|  |  | Hydrophilic BL | 2.510 | 0.465 | 1.586 – 3.435 | 1.0002 | 13870.6 | 13268.6 | < 0.001 *** |
|  |  | Mesic BL | -0.129 | 0.394 | -0.907 – 0.650 | 1.0006 | 13580.9 | 12762.9 | 0.734 |
|  | Var.pars. | Residual SD | 0.969 | 0.126 | 0.761 – 1.253 | 1.0001 | 14859.1 | 12855.7 | – |
| Temp. | Fixed | Intercept | 3.562 | 0.335 | 2.898 – 4.217 | 1.0004 | 13822.1 | 11956.8 | < 0.001 *** |
|  |  | Hydrophilic BL | 1.805 | 0.464 | 0.892 – 2.730 | 1.0001 | 13936.5 | 13376.7 | < 0.001 *** |
|  |  | Mesic BL | 2.740 | 0.422 | 1.907 – 3.571 | 1.0004 | 14091.0 | 13493.0 | < 0.001 *** |
|  | Var.pars. | Residual SD | 0.949 | 0.132 | 0.736 – 1.248 | 1.0002 | 14814.0 | 12765.2 | – |
| Light | Fixed | Intercept | 6.887 | 0.282 | 6.334 – 7.451 | 1.0001 | 14916.0 | 12968.7 | < 0.001 *** |
|  |  | Hydrophilic BL | -0.226 | 0.401 | -1.010 – 0.569 | 1.0004 | 15478.2 | 13713.0 | 0.556 |
|  |  | Mesic BL | -0.943 | 0.369 | -1.666 – -0.209 | 1.0002 | 15345.1 | 14032.2 | 0.014 * |
|  | Var.pars. | Residual SD | 0.879 | 0.116 | 0.687 – 1.137 | 1.0003 | 17224.4 | 14070.0 | – |

**Table S7.** Results of contrast tests of differences in Ellenberg-Tichý indicator values. Shown are the posterior means of differences with their MCMC standard error, 95 % credible interval, and a Bayesian pseudo-p-value *P_Bayes_* (see Notes S1.4). Symbols indicate parameters credibly different from zero on the 0.1 (.), 0.05 (*), 0.01 (**) and 0.001 level (***).

| Variable | Group 1 | Group 2 | Estimate | Est. Error | 95% CI | $P_{Bayes}$ |
| --- | --- | --- | --- | --- | --- | --- |
| Moisture | mesic | hydrophilic | -2.639 | 0.424 | -3.475 – -1.796 | < 0.001 *** |
|  | conifer | mesic | -0.129 | 0.394 | -0.907 – 0.650 | 0.734 |
|  | conifer | hydrophilic | 2.510 | 0.465 | 1.586 – 3.435 | < 0.001 *** |
| Temperature | mesic | hydrophilic | 0.934 | 0.408 | 0.124 – 1.741 | 0.025 * |
|  | conifer | mesic | 2.740 | 0.422 | 1.907 – 3.571 | < 0.001 *** |
|  | conifer | hydrophilic | 1.805 | 0.464 | 0.892 – 2.730 | < 0.001 *** |
| Light | mesic | hydrophilic | -0.716 | 0.367 | -1.446 – 0.009 | 0.053 . |
|  | conifer | mesic | -0.943 | 0.369 | -1.666 – -0.209 | 0.014 * |
|  | conifer | hydrophilic | -0.226 | 0.401 | -1.010 – 0.569 | 0.556 |

**Table S8.** Derived quantities from the model fits for all models in Table S2. Shown are the posterior mean with its 95 % credible intervals and a Bayesian pseudo-p-value *P_Bayes_* (see Notes S1.4) for the total correlation between log(−*P*50) and log(*K*s), the standardized major axis intercept and slope for the relationship across species, the slope for the relationship within species as well as the difference between the across-and within-species slope. Symbols indicate parameters credibly different from zero on the 0.1 (.), 0.05 (*), 0.01 (**) and 0.001 level (***).

| Model | Parameter | Estimate | 95% CI | | $P_{Bayes}$ |
| --- | --- | --- | --- | --- | --- |
| Pooled | Corr. total | -0.370 | -0.546 | -0.175 | < 0.001 *** |
|  | Intercept (across) | 2.365 | 1.838 | 2.979 | < 0.001 *** |
|  | Slope (across) | -1.848 | -2.347 | -1.427 | < 0.001 *** |
|  | Slope (within) | -4.692 | -5.442 | -4.013 | 0.001 ** |
|  | Slope difference | -2.843 | -3.730 | -1.996 | 0.001 ** |
| Hydrophilic broadleaf | Corr. total | -0.058 | -0.367 | 0.243 | 0.721 |
|  | Intercept (across) | 1.308 | -2.841 | 4.805 | 0.523 |
|  | Slope (across) | -0.980 | -5.796 | 4.712 | 0.648 |
|  | Slope (within) | -0.117 | -7.988 | 8.095 | 0.978 |
|  | Slope difference | 0.864 | -10.216 | 11.264 | 0.998 |
| Mesic broadleaf | Corr. total | -0.490 | -0.677 | -0.270 | < 0.001 *** |
|  | Intercept (across) | 2.429 | 1.779 | 3.260 | < 0.001 *** |
|  | Slope (across) | -1.343 | -1.957 | -0.859 | < 0.001 *** |
|  | Slope (within) | -5.622 | -7.096 | -4.430 | 0.009 ** |
|  | Slope difference | -4.279 | -5.889 | -2.879 | 0.009 ** |
| Conifer | Corr. total | -0.252 | -0.482 | 0.001 | 0.051 . |
|  | Intercept (across) | 0.423 | -1.993 | 1.478 | 0.260 |
|  | Slope (across) | -0.704 | -1.496 | 1.121 | 0.231 |
|  | Slope (within) | -3.526 | -4.477 | -2.735 | 0.005 ** |
|  | Slope difference | -2.822 | -4.844 | -1.605 | 0.005 ** |

